# Polo-Like Kinase 1 phosphorylation tunes the functional viscoelastic properties of the centrosome scaffold

**DOI:** 10.1101/2024.08.29.610374

**Authors:** Matthew Amato, June Ho Hwang, Manolo U. Rios, Nicole E. Familiari, Michael K. Rosen, Jeffrey B. Woodruff

## Abstract

Centrosomes are membranelles organelles containing centrioles encapsulated by pericentriolar material (PCM). PCM nucleates microtubules that help position and segregate chromosomes during mitosis, yet how PCM resists microtubule-mediated forces is poorly understood at the material level. Here, we show that PLK-1 phosphorylation of SPD-5 tunes the dynamics and material properties of the PCM scaffold in *C. elegans* embryo*s.* Microrheology of reconstituted PCM condensates reveals that PLK-1 phosphorylation decreases SPD-5 dynamics and increases condensate viscoelasticity. Similarly, in embryos, phospho-mimetic SPD-5 is less dynamic than wild-type SPD-5, which itself is less dynamic than phospho-null SPD-5. PCM built with phospho-null SPD-5 is smaller than normal, but its assembly can be partially rescued by reducing microtubule-dependent forces. The same is true for PCM built with phospho-mimetic SPD-5, yet the underlying causes are distinct: under force, phospho-null SPD-5 fails to assemble, while phospho-mimetic SPD-5 forms hyper-stable foci that fail to cohere into a uniform, spherical mass. Both mutants have defects with chromosome segregation and viability. Thus, tuning of SPD-5 phosphorylation optimizes PCM material properties to achieve correct PCM size, integrity, and function. Our results demonstrate how regulated chemical modification of a scaffolding protein modulates the material properties and function of a membraneless organelle.

## 1.0 INTRODUCTION

Mesoscale material properties like viscosity, elasticity, and strength emerge from the collective interactions between molecules. This is well understood for associative polymers solutions that undergo gelation or phase separation (Cohen, 1982; Flory, 1953). For example, chain length, valence, strength of interactions, and solubility are all well characterized to influence the material properties of engineered and biological polymer-based materials (Alshareedah et al., 2024; Ferry, 1980; Harmon et al., 2017). While this framework has been applied to understand subcellular structures, such as membraneless organelles (Banani et al., 2017; Lyon et al., 2021), the relationship between material properties and function of such structures is incompletely understood.

The centrosome represents an ideal model to investigate how material properties influence the function of membraneless organelles. The centrosome is a force-bearing structure composed of barrel-shaped centrioles that organize an amorphous proteinaceous matrix known as the pericentriolar material (PCM). PCM-nucleated microtubules help form the mitotic spindle, which segregates mitotic chromosomes during cell division (Gomes Pereira et al., 2021; Woodruff et al., 2014). PCM also nucleates astral microtubules that connect to cortically anchored motors, which generate force to position the mitotic spindle (Laan et al., 2012). If PCM integrity is disrupted, microtubule-based pulling forces can fragment it, leading to multipolar spindles and genomic instability (Maiato and Logarinho, 2014; Oshimori et al., 2006). These observations suggest that the material properties of the PCM are functionally important.

Understanding the molecular interactions within PCM could be informative for determining functionally important material properties. Upon entry into mitosis, PCM rapidly expands by accumulating scaffold proteins that then recruit client proteins needed for microtubule aster formation (Gomes Pereira et al., 2021). Major scaffold proteins have been identified in nematodes (SPD-5), insects (Cnn), and vertebrates (CDK5RAP2). These all contain numerous coiled-coil domains interspersed with disordered linkers (Hamill et al., 2002; Megraw et al., 1999) Polo-family kinase phosphorylation potentiates PCM maturation by promoting the self-association of these scaffold proteins (Conduit et al., 2014; Feng et al., 2017; Haren et al., 2009; Rios et al., 2024; Rios et al., 2025; Woodruff et al., 2015) In *C. elegans,* Polo-like Kinase (PLK-1) phosphorylation of SPD-5 at four sites (S530, S626, S653, S658) is important for SPD-5 multimerization and overall PCM maturation (Woodruff et al., 2015). Optical nano-rheology showed that *C. elegans* PCM can flow under shear stress, but only during its disassembly phase or when PLK-1 is inhibited (Mittasch et al., 2020). This result suggests that the material properties of the PCM are dynamic and are subject to cell cycle regulation by PLK-1. It is unclear if PLK-1 directly regulates the material properties of the PCM and whether properly tuned material properties are important for PCM function.

Here, we used cell-based and *ex vivo* biophysical assays, biochemical reconstitution, and microrheology to characterize how PLK-1 regulates the material properties of the PCM in *C. elegans*. We demonstrate that PLK-1 phosphorylation increases the viscoelastic moduli of reconstituted SPD-5 scaffolds in vitro. In embryos, we show that SPD-5 can transition between dynamically and stably associated states based on its phosphorylation status. A balance of these two states tunes the material properties of the PCM to optimize its mechanical integrity and ability to segregate chromosomes during mitosis.

## 2.0 Results

### 2.1 PLK-1 phosphorylation changes the dynamics and material properties of reconstituted PCM scaffolds

To investigate how PLK-1 phosphorylation changes the material properties of the PCM, we biochemically reconstituted PCM scaffold assembly *in vitro.* Previous reconstitution of micron-scale SPD-5 condensates required the use of macromolecular crowding agents, which dampened regulation by PLK-1 (Woodruff et al., 2017). More importantly, the presence of crowding agents has been shown to artificially affect the elasticity and viscosity of biomolecular condensates (Andre et al., 2023; Kaur et al., 2019). To circumvent these complications, we discovered conditions that permit SPD-5 condensation without using a macromolecular crowding agent. We found that SPD-5 in combination with its client protein TPXL-1 condenses into micron-scale droplets at near-physiological salt concentrations (75 mM KCl; Figure 1A, S1A,B). For the rest of this study, we refer to these reconstituted assemblies as “PCM condensates.”

**Figure 1:**
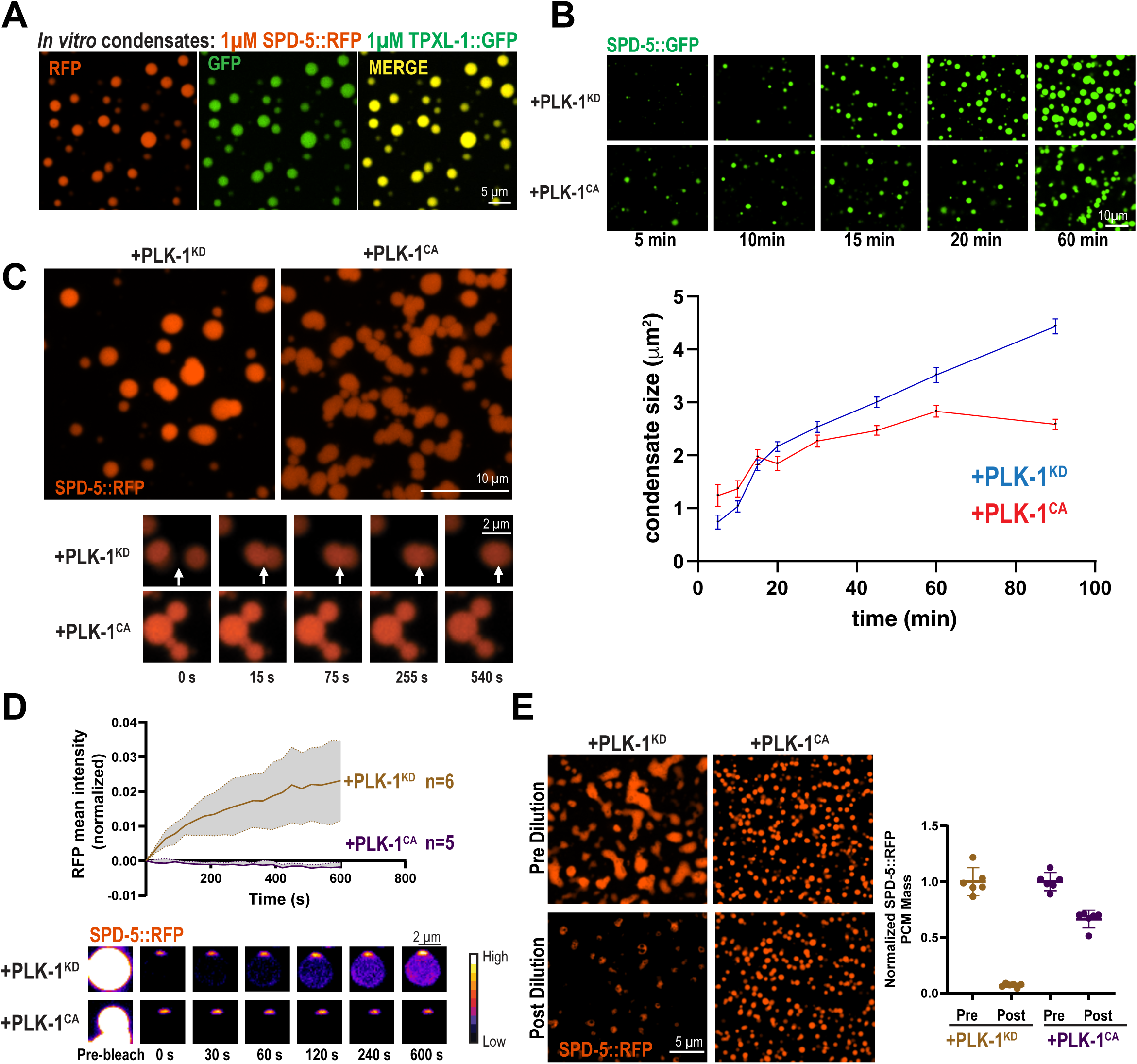
PLK-1 phosphorylation changes the dynamics and material properties of reconstituted PCM scaffolds. a. Reconstitution of PCM condensates using purified SPD-5, TPXL-1, and no molecular crowding agent. Proteins were assembled for 15 min in buffer (75 mM KCl, 4 mM Hepes pH 7.4, 10 mM DTT, 0.4 mM ATP, 1 mM MgCl_2_) and imaged in a 96-well glass bottom plate using confocal fluorescence microscopy. b. *In vitro* PCM condensate assembly assay over time. Top, representative images. Bottom, quantification of average condensate size over time (mean +/− 95% C.I.; n>100 condensates from 3 replicates). c. Samples were prepared as in (A) with 0.5 µM kinase dead PLK-1(KD) or constitutively active PLK-1(CA) and imaged after 30 min. Top panel, static images. Bottom panel, timelapse imaging of condensates. Arrows indicate the site of condensate fusion. d. Fluorescence recovery after photobleaching of SPD-5/TPXL-1 condensates. Top, SPD-5::RFP intensity was measured and normalized (mean +/− 95% C.I.; PLK-1(KD) n = 6, PLK-1(CA) n = 6 condensates). Bottom, images of bleached condensates. e. Dilution assay of reconstituted PCM. 1 µM SPD-5::RFP, 1 µM TPXL-1::GFP was incubated in buffer (50 mM KCl, 4 mM Hepes pH 7.4, 10 mM DTT, 0.4 mM ATP, 1 mM MgCl_2_) with 0.5 µM PLK-1 (KD) or 0.5 µM PLK-1(CA) for 1 hr and imaged. Samples were then diluted 4.3X into extrusion buffer (150mM KCl, 25mM Hepes pH.7.4) and imaged after 1 hr (left panels). Right, quantification of SPD-5::RFP integrated density before and after dilution (mean +/− 95% C.I.; PLK-1(KD) n=6 images, PLK-1 (CA) n=6 images with >100 condensates).

Time-lapse imaging revealed that, after mixing of SPD-5, TPXL-1, and kinase dead (KD) PLK-1, PCM condensates steadily grew. Incubation with constitutively active (CA) PLK-1 initially enhanced PCM condensate assembly compared with the KD control (<10 min) (Figure 1B). Thus, PLK-1 activity, and not simply binding, promotes condensate assembly, recapitulating PLK-1-mediated potentiation of PCM assembly seen in vivo (Decker et al., 2011; Woodruff et al., 2015). At later time points, however, PLK-1 activity inhibited PCM condensate growth, suggestive of kinetic arrest (Linsenmeier et al., 2022; Ranganathan and Shakhnovich, 2020). Furthermore, PCM condensates incubated with PLK-1(CA) appeared clustered, reminiscent of a colloidal suspension, whereas condensates incubated with PLK-1(KD) were dispersed (Figure 1B,C).

Time-lapse imaging revealed that PCM condensates with PLK-1(KD) could fuse, while condensates incubated with PLK-1(CA) could not fuse and remained clustered after 540 s (Figure 1C). Molecular rearrangement underlies droplet coalescence in soft matter materials; therefore, we hypothesized that PLK-1 activity decreases the dynamics of PCM condensates. Indeed, condensates with PLK-1(KD) recovered after photobleaching, albeit at a low level, whereas condensates with PLK-1(CA) showed no recovery (Figure 1D, S1B). As an orthogonal approach, we measured persistence of SPD-5::RFP in condensates after 4.3-fold dilution into a higher salt buffer (150mM KCl, 25mM Hepes pH 7.4). Unphosphorylated condensates lost considerable mass upon dilution, whereas phosphorylated condensates largely maintained their mass (Figure 1E).

To confirm that dynamic changes conferred by PLK-1 activity were due to the phosphorylation state of SPD-5 and not TPXL-1, we used phospho-mimetic SPD-5, which contains serine to glutamic acid substitutions at 4 PLK-1 phosphorylation sites important for PCM assembly (S530E, S627E, S653E, S658E; SPD-5(4E)) (Woodruff et al., 2015). SPD-5(4E) condensates morphologically and dynamically recapitulated SPD-5(WT) condensates treated with PLK-1(CA); both assembled smaller drops unable to fuse that were resistant to dilution (Fig S1C, D). Furthermore, PLK-1 did not reduce TPXL-1 dynamics but rather increased them (Figure S1E). We conclude that phosphorylation of SPD-5 directly decreases the dynamics of reconstituted PCM scaffolds and their ability to fuse.

### 2.2 Phosphorylation increases the viscoelasticity of reconstituted PCM scaffolds

To quantify how phosphorylation changes the material properties of reconstituted PCM, we used 100 nm beads to perform video particle tracking microrheology (VPT) of SPD-5/TPXL-1 condensates incubated with PLK-1(CA) or PLK-1(KD)(Figure 2A). To minimize effects from confinement, only beads located in the condensate center were analyzed (see methods). Phosphorylation reduced the mean squared displacement (MSD) of beads inside the condensates (n>17,000 trajectories; Figure 2B). Using the Generalized Stokes-Einstein Relation, we expressed MSD in terms of the viscoelastic modulus, which consists of viscous (G”) and elastic (G’) moduli as functions of frequency (ω) (Figure 2C). Phosphorylation shifted the two moduli upward, indicating a transition to higher viscoelasticity (Figure 2C), and shifted the first crossover point (where G’ = G”) to the left, indicating emergence of elastic (solid-like) behavior at lower frequencies. In PLK-1(CA) condensates, G’ and G’’ appeared to approach a second crossover point at a higher frequency not accessible by our VPT analysis. We speculate that phosphorylation could qualitatively change the rheological behaviors of SPD-5 condensates in addition to simply quantitatively increasing their viscoelasticity (see Discussion). Next, we derived viscosity (η) as a function of frequency using the relation η=G”/ω. Phosphorylation increased the condensate viscosity at all frequencies (Figure 2D). The estimated zero-shear viscosity (resting viscosity) of the condensates differed more than 8-fold: 31.8 Pa*s in the presence of PLK-1(CA) vs. 3.8 Pa*s in the presence of PLK-1(KD).

**Figure 2:**
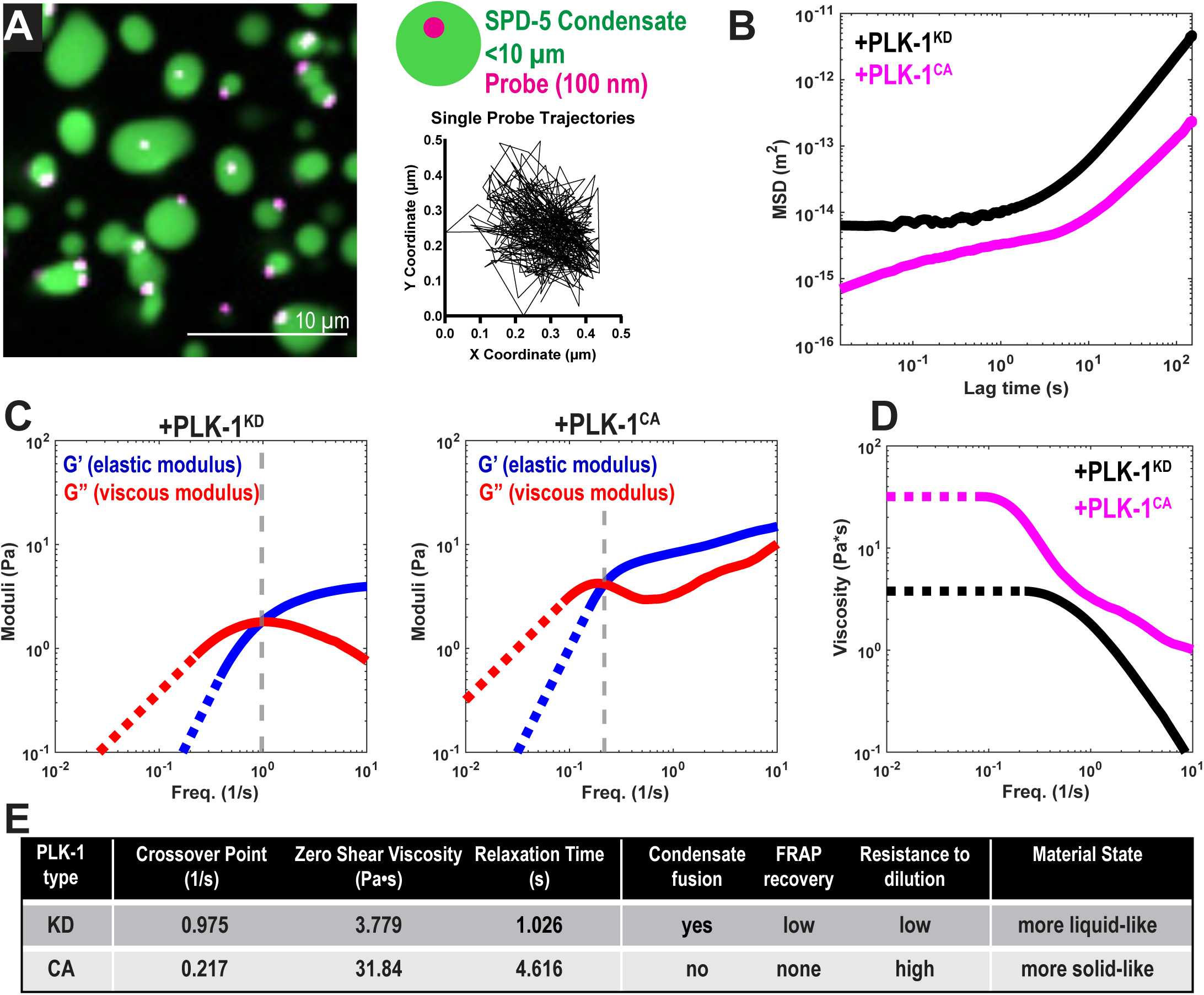
Phosphorylation increases the viscoelasticity of reconstituted PCM scaffolds. a. Schematic of Video Particle Tracking (VPT) technique. 100 nm-diameter carboxylate-modified fluorescent beads were encapsulated in reconstituted PCM condensates. Left, representative spinning-disc confocal fluorescence image. Top right, diagram of condensate with bead; not to scale. Bottom right, representative trajectory of a single bead within a condensate. b. Plot for averaged mean squared displacement (MSD) of beads encapsulated in reconstituted PCM condensates incubated with 0.2 µM PLK-1(KD) or PLK-1(CA). c. Viscoelastic moduli of condensates determined from VPT. Moduli were calculated using the generalized Stokes-Einstein relation fit to a maxwell fluid from the averaged MSD from 0.015 to 100 s. For G’ and G”, solid lines indicate measured values, dashed lines indicate extrapolated values. Dashed grey line indicates crossover frequency. d. Viscosity was calculated from averaged MSD from 0.015 to 100 s. The mean value of the plateau at the low frequency was used to estimate the zero-shear viscosity (η_0_). Solid lines indicate measured values, dashed lines indicate extrapolated values. e. Summary of in vitro rheological and dynamics measurements.

The response of a viscoelastic material is dominated by elastic contributions on short time scales and viscous contributions on longer time scales. The terminal relaxation time characterizes the timescale when a material transitions from elastic to terminally viscous behavior under applied stress and is defined as the inverse of the frequency at the first G’-G” crossover point. Longer terminal relaxation times are associated with more elastic, solid-like states. Phosphorylation increased the terminal relaxation time of SPD-5/TPXL-1 condensates (4.6 s (CA) vs.1.03 s (KD); Figure 2C). Our microrheology data, combined with FRAP and fusion assays, indicate that PLK-1 phosphorylation increases the viscoelasticity of the PCM condensates, which leads to more solid-like PCM at functionally relevant time scales. Furthermore, our data demonstrate that the dynamics of SPD-5 correlate with viscoelasticity of PCM condensates (Figure 2E). We conclude that phosphorylation-dependent tuning of SPD-5 dynamics determines the overall viscoelastic behavior of PCM condensates.

### 2.3 The phosphorylation state of SPD-5 affects its dynamics in metaphase-arrested embryos

We next investigated how phosphorylation impacts PCM material properties and function in embryos. It is technically challenging to introduce beads into *C. elegans* PCM and perform VPT. However, our in vitro system showed a strong correlation between phosphorylation-induced changes in dynamics and material properties of SPD-5 condensates. Therefore, we sought to infer PCM material properties in embryos by measuring SPD-5 dynamics. We generated embryos that express GFP-tagged, RNAi-resistant transgenes of *spd-5* from an ectopic locus in the genome (Woodruff et al., 2015). We mutated the four residues (S530, S627, S653, S658) known to be phosphorylated by PLK-1 and important for PCM assembly to alanine (4A; phospho-null) or glutamic acid (4E; phospho-mimetic) (Figure 3A). Importantly, this system allows us to test the effects of mutant protein dosage: without RNAi, both mutant and wild-type SPD-5 co-exist, while upon RNAi depletion of endogenous *spd-5*, only the mutant version is expressed (Wueseke et al., 2016).

**Figure 3:**
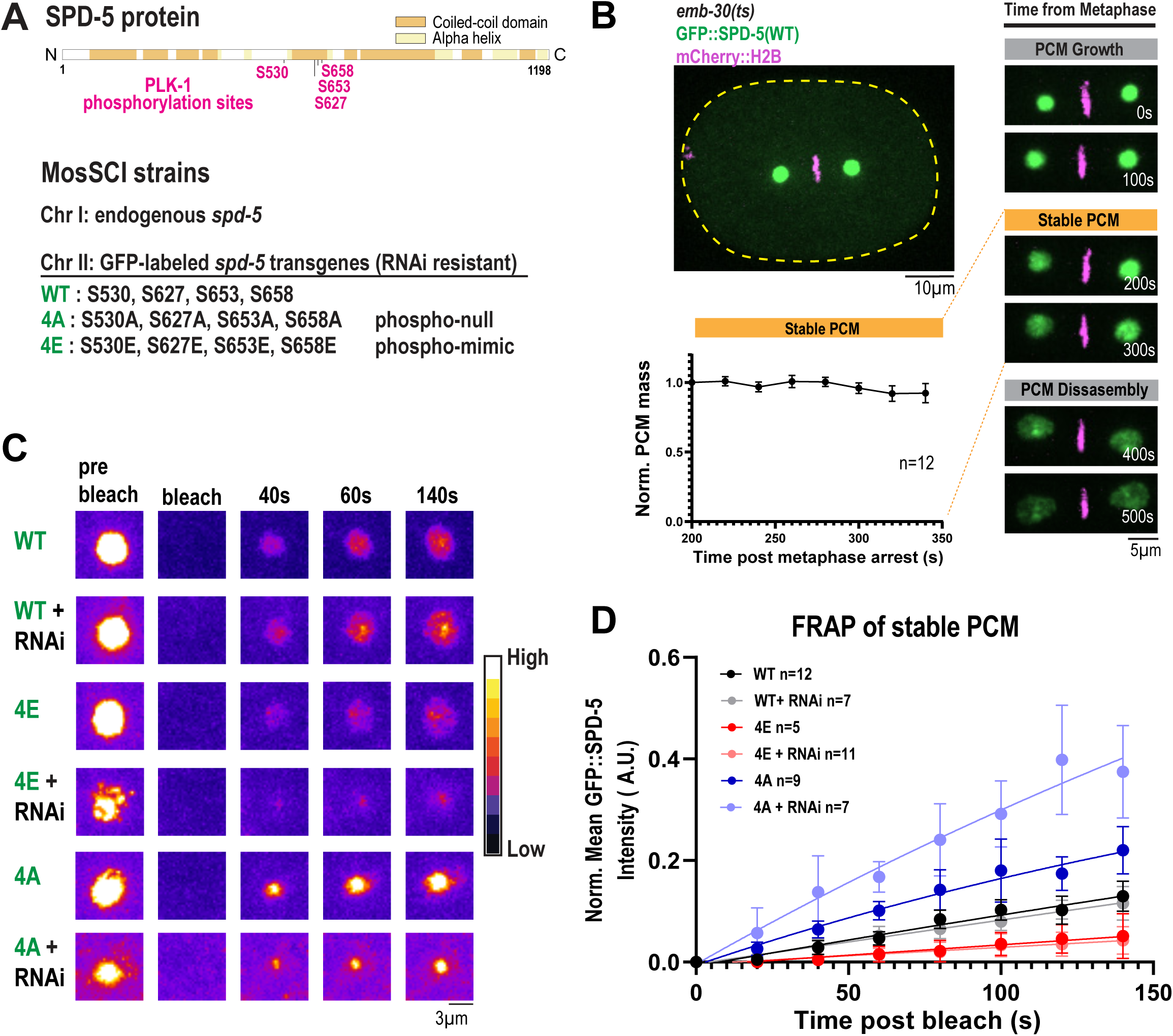
The phosphorylation state of SPD-5 affects its dynamics in metaphase-arrested embryos. a. Top, diagram of key PLK-1 phosphorylation sites in SPD-5. Bottom, design of *gfp::spd-5* transgenes expressed at the Mos locus on chromosome II. b. Top, representative image of metaphase arrest in one-cell *emb-30(ts)* embryos expressing GFP::SPD-5(WT) and mCherry::H2B. Right, characterization of PCM size during metaphase arrest. Bottom, normalized integrated density of GFP::SPD-5 200-340 s after metaphase arrest (mean +/− 95% C.I.; n= 12 embryos). c. Representative images for fluorescence recovery after photobleaching (FRAP) of PCM in metaphase-arrested embryos. PCM was photobleached at 200 s post metaphase arrest in one-cell embryos expressing *gfp::spd-5* transgenes with and without RNAi against endogenous *spd-5*. d. Quantification of (C). GFP mean intensity was measured every 20 s until 140 s and normalized (mean +/− 95% C.I.; GFP::SPD-5(WT) n= 12, GFP::SPD-5(WT) + *spd-5(RNAi) n= 7,* GFP::SPD-5(4E) n=8, GFP::SPD-5(4E) + *spd-5(RNAi)* n=11, GFP::SPD-5(4A) n=9, GFP::SPD-5(4A) + *spd-5(RNAi)* n=7 centrosomes). Curves are fit to a one-phase nonlinear regression model.

We assayed how the phosphorylation state of SPD-5 affects PCM scaffold dynamics by performing fluorescence recovery after photobleaching (FRAP). To decouple signal recovery contributed by replacement of existing material (dynamics) from incorporation of new material from the cytosol (growth), we arrested cells in metaphase when PCM stops growing (Laos et al., 2015). This is achieved by expressing a temperature sensitive allele of *emb-30,* which encodes a subunit of the Anaphase Promoting Complex (Furuta et al., 2000). After shifting embryos to 25°C for 15 min, confocal imaging verified that embryos were arrested in metaphase and that PCM was most stable 200-340 s post arrest. After 400 s following arrest, PCM disassembled, indicating escape from arrest (Figure 3B). Therefore, we restricted our analysis to 200-340 s post arrest.

In the presence of endogenous SPD-5 (no RNAi), GFP::SPD-5(4A) was the most dynamic, recovering up to ∼20%, whereas GFP::SPD-5(WT) and GFP::SPD-5(4E) recovered at ∼10% and ∼5% respectively (Figure 3C,D). To test the effects of the mutant proteins alone, we depleted endogenous SPD-5 using RNAi. Again, GFP::SPD-5(4A) was the most dynamic, followed by WT then 4E (Figure 3C,D). RNAi treatment increased the dynamics of SPD-5(4A), implying that it is partly stabilized by the presence of wild-type protein (Figure 3D). On the other hand, RNAi treatment did not affect the dynamics of the WT or 4E mutant. We noticed that PCM made with the SPD-5(4E) was highly irregular, comprising an aggregate of several foci, rather than a cohesive, spherical assembly; the nature of this behavior is not clear but could be a consequence of kinetic arrest (see Discussion). These data indicate that a small, dynamic population of SPD-5 exists in metaphase PCM. Furthermore, phosphorylation decreases the dynamics of SPD-5 within PCM, consistent with our in vitro results.

### 2.4 SPD-5 dynamics change with the cell cycle

Our data indicate that PCM scaffold dynamics are affected by the phosphorylation state of SPD-5. Given that PLK-1 activity changes during the cell cycle (Golsteyn et al., 1995; Mittasch et al., 2020), we hypothesized that PCM scaffold dynamics are tuned during the cell cycle. Due to the fast movement and growth of PCM, it is difficult to measure dynamics using FRAP. Therefore, we used laser-induced extrusion to expel PCM from the cell and measured its persistence, similar to our *in vitro* dilution assay (Figure 1E). We performed extrusion by coating *C. elegans* embryos with calcofluor white, a fluorescent dye that binds chitin and absorbs blue light, then using a short, focused pulse of 405 nm laser light to create precise punctures in the embryo cortex. Application of light pressure forced PCM out of the embryo into a dilute, controlled medium (Figure 4A). To ensure that the extrusion process itself does not damage the integrity of PCM, we first extruded PCM (labeled with GFP::SPD-5) into low salt buffer (25 mM HEPES, 0 KCl). PCM remained mostly stable for >1 hr (Figure 4B); thus, the wild-type PCM scaffold is not damaged by extrusion. These results demonstrate that PCM is intrinsically stable and does not require cytoplasmic factors for its maintenance in low salt conditions.

**Figure 4:**
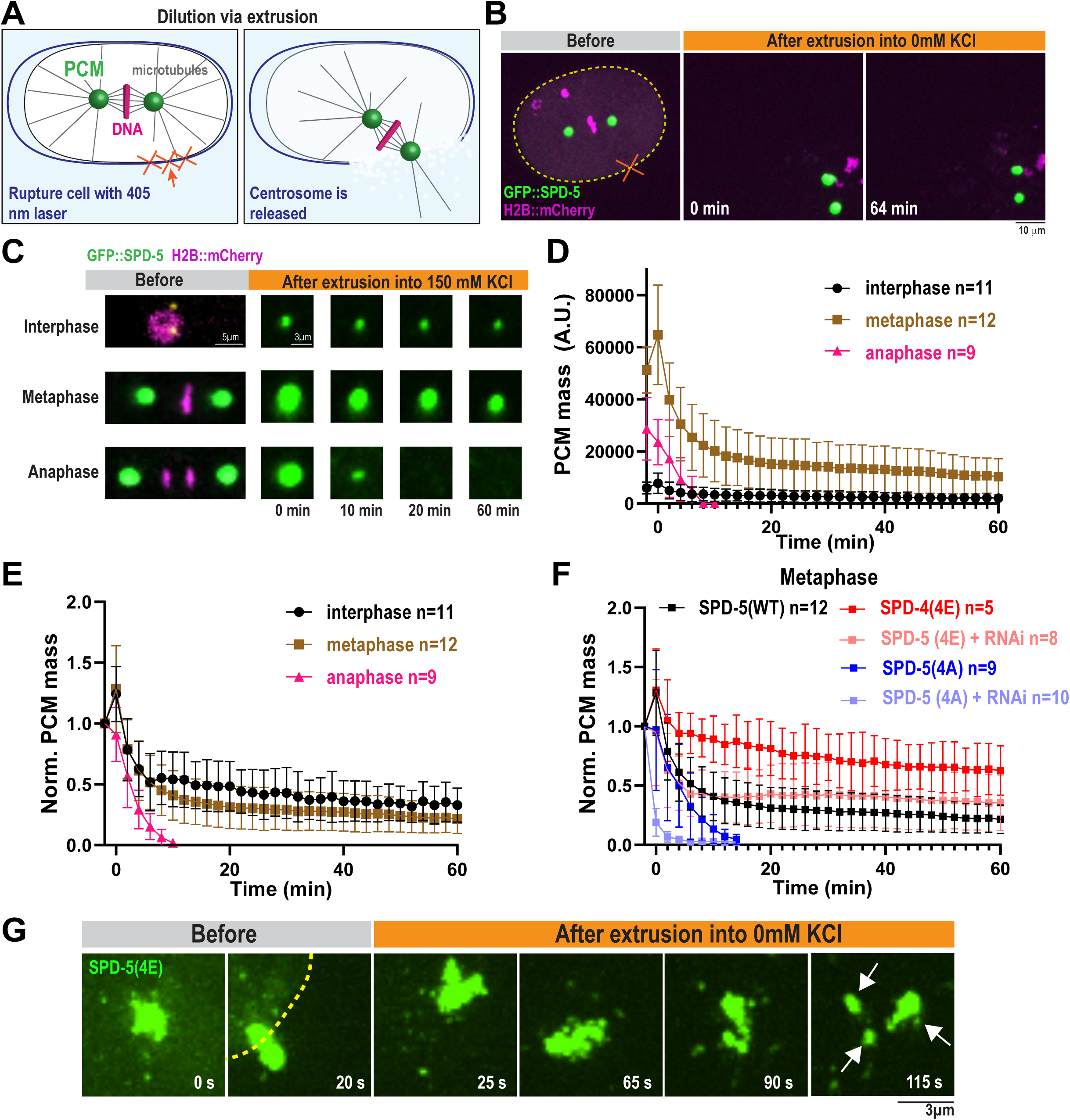
SPD-5 dynamics change with the cell cycle. a. Diagram of centrosome release and dilution via extrusion. *C. elegans* embryos were coated with calcofluor white and eggshells were ruptured with a 405 nm laser. b. Representative images of centrosomes (labeled with GFP::SPD-5) extruded from a one-cell embryo at metaphase into low salt buffer (0mM KCl, 25mM HEPES pH 7.4). DNA is labeled with H2B::mCherry. c. Representative images of centrosomes extruded from one-cell embryos in different cell cycle stages into high salt buffer (150mM KCl, 25mM HEPES pH 7.4). d. Quantification of integrated density of GFP::SPD-5 of extruded centrosomes imaged every 2 minutes for 1 hr (mean +/− 95% C.I.; interphase n=11, metaphase n=12, anaphase n=9 centrosomes). e. Normalized data from D. f. Normalized GFP integrated density of extruded centrosomes from metaphase embryos expressing *gfp::spd-5 transgenes* with and without *spd-5(RNAi)*(mean +/− 95% C.I.; GFP::SPD-5(WT) n= 12, GFP::SPD-5(4E) n=5, GFP::SPD-5(4E) + *spd-5(RNAi)* n=8, GFP::SPD-5(4A) n=9, GFP::SPD-5(4A) + *spd-5(RNAi)* n=10 centrosomes). g. Representative images of extruded centrosomes in one-cell metaphase embryos expressing *gfp::spd-5(4E)* treated with *spd-5(RNAi)* into low salt buffer (0mM KCl, 25mM Hepes pH 7.4).

Extrusion into a high salt buffer (25 mM HEPES, 150 mM KCl) caused dissipation of GFP::SPD-5 signal in a cell-cycle specific manner (Figure 4C,D). In extruded interphase and metaphase cells, GFP integrated density rapidly declined initially and then plateaued for up to 60 min, whereas, in anaphase, GFP integrated density monotonically decreased and was no longer detected after ∼10 min (Figure 4C-E). The total amount of GFP signal remaining was higher in metaphase vs. interphase. However, normalization of the signal revealed that departure rates were similar between metaphase and interphase (Figure 4E). Our results indicate that PCM scaffold dynamics are subject to cell cycle-dependent regulation: dynamics are relatively low in interphase and metaphase, then increase dramatically in anaphase, which could facilitate PCM disassembly. Furthermore, our data show the utility of extrusion to probe the dynamics of growing or moving organelles.

We next wondered how disrupted phospho-regulation affects PCM persistence following extrusion. We chose to focus on metaphase cells, when PCM is at its largest and perturbations in stability, size, and integrity would be most prominent. In the presence of endogenous SPD-5, GFP::SPD-5(4A) signal rapidly disappeared after extrusion (Figure 4F), reminiscent of wild-type anaphase PCM. On the other hand, GFP::SPD-5(4E) strongly persisted, much longer than GFP::SPD-5(WT) (Figure 4F). Considering that the phosphorylation state of SPD-5 affected the dynamics of extruded PCM, we wondered if PP2A, a PCM-resident phosphatase, could dephosphorylate PCM after extrusion and affect its dynamics. Extrusion of metaphase PCM into a high salt buffer with 10 µM LB-100, which inhibits PP2A (Magescas et al., 2019), showed higher persistence compared to the control (Figure S2). This result suggests that PCM-associated PP2A can dephosphorylate SPD-5 after extrusion and facilitate its departure. We conclude that phosphorylation of SPD-5 increases the persistence of the PCM scaffold following extreme dilution upon extrusion. These data support that phosphorylation decreases the rate of SPD-5 dissociation from PCM, consistent with our in vitro results.

In the absence of endogenous SPD-5 (via *spd-5(RNAi)*), GFP::SPD-5(4A) signal was lost even quicker (Figure 4F), again suggesting that wild-type SPD-5 can help stabilize unphosphorylated SPD-5, as we saw in our FRAP assay. Unexpectedly, SPD-5(4E) integrated density rapidly decreased initially and then stabilized over time; this pattern was markedly different from the no RNAi condition. Higher resolution imaging revealed that PCM made exclusively from SPD-5(4E) fragmented immediately following extrusion, whereas wild-type PCM remained intact (Figure 4G). These 4E fragments remained stable in fluorescence intensity but often drifted out of the imaging plane, thus giving the appearance of lower overall GFP integrated density. We conclude that PCM built with hyper-phosphorylated SPD-5 has compromised mechanical integrity at the meso-scale: SPD-5(4E) can form small assemblies that are highly stable but poorly connected to other assemblies. This lack of overall cohesiveness could explain the irregular shape of PCM in *spd-5(4E)* embryos.

### 2.5 The phosphorylation state of SPD-5 tunes the mechanical integrity of the PCM

We then examined how the viscoelastic properties of PCM contribute to its mechanical integrity, specifically its ability to withstand external forces without fracturing or deforming. PCM must resist microtubule pulling forces, and defects in PCM scaffold assembly can cause premature, force-driven disassembly of PCM (Rios et al., 2024). Our in vitro experiments indicated the viscoelastic properties of the SPD-5 scaffold correlate with SPD-5 dynamics, and that both are regulated by PLK-1 phosphorylation. Given that SPD-5(4E) PCM without endogenous SPD-5 is structurally unsound when extruded from embryos, we hypothesized that this PCM also suffers an assembly defect that could be rescued by removing microtubule pulling forces by depolymerizing microtubules with nocodazole.

Confocal imaging revealed that in the absence of endogenous SPD-5, PCM built from GFP::SPD-5(4E) accumulated less mass relative to WT during mitosis (Figure S3A). PCM was also smaller when SPD-5(4E) was expressed in the presence of endogenous SPD-5 (Figure S3A). These changes in size were not due to differences in expression levels or cell cycle lengths (Figure S3B,C). PCM built solely from SPD-5(4E) appeared irregular and broken, in contrast to uniform, spherical wild-type PCM (Figure 5A). To test if this is a material defect, we eliminated pulling forces by treating embryos with nocodazole. Application of 40 μM nocodazole fully rescued the morphology of PCM in *spd-5(4E)* embryos and partially rescued PCM mass (1.82-fold increase; p < 0.005), while PCM mass in *spd-5(WT)* embryos was not affected (p > 0.9)(Figure 5A,B). Thus, *spd-5(4E)* embryos have both a force-dependent and -independent deficiency in PCM assembly. We conclude that hyper-phosphorylation of SPD-5 impairs its ability to build a cohesive, full-sized scaffold.

**Figure 5:**
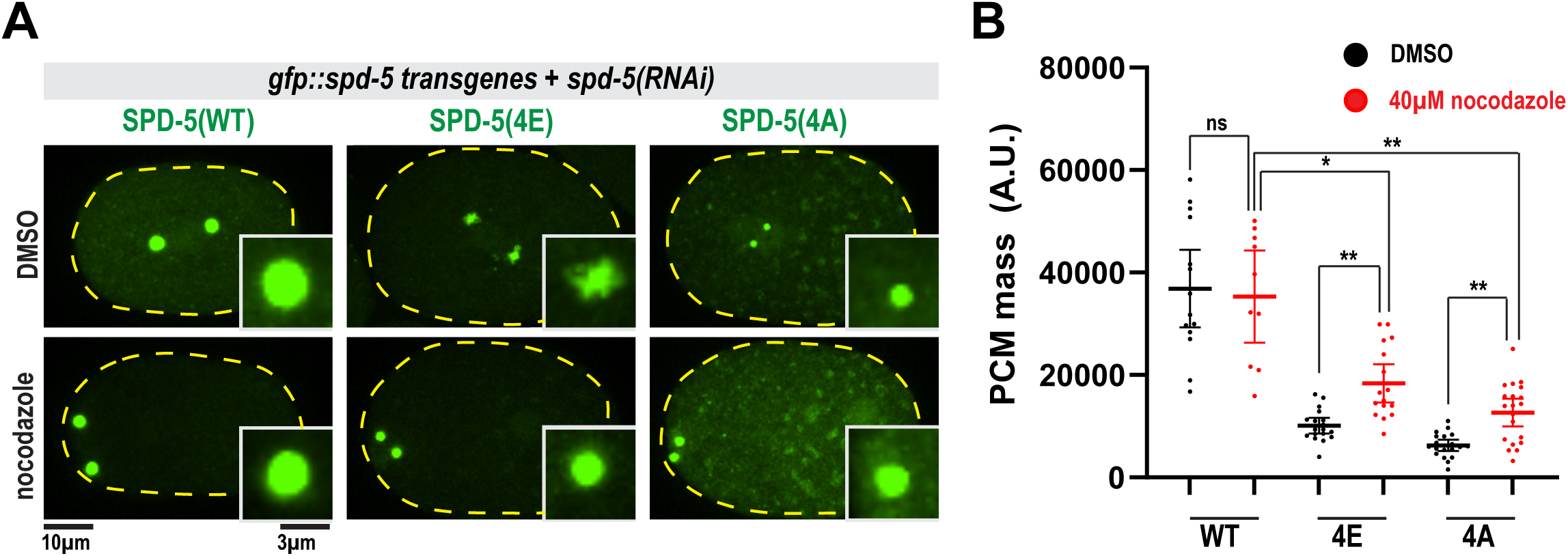
The phosphorylation state of SPD-5 is tuned to maximize the mechanical integrity of the PCM. a. Transgenic worms were treated with *spd-5* RNAi to deplete endogenous SPD-5. Left, one-cell embryos expressing *gfp::spd-5 transgenes* treated with 2% DMSO or 40 µM nocodazole and imaged at nuclear envelope breakdown. b. Quantification GFP integrated density at PCM (mean +/− 95% C.I.; WT + DMSO n = 14, WT+ nocodazole n=10, 4E + DMSO n=18, 4E + nocodazole n=16, 4A + DMSO n=20, 4A + nocodazole n=20; p values from Brown-Forsythe and Welch ANOVA tests, followed by a Dunnet’s multiple comparisons test).

Embryos expressing only SPD-5(4A) failed to build full-sized PCM, consistent with previous studies (Woodruff et al., 2015). This phenotype was previously interpreted exclusively as a defect in PCM assembly. However, treatment with 40 μM nocodazole partially rescued PCM scaffold mass in these embryos (2-fold; p < 0.005)(Figure 5A,B). Thus, similar to SPD-5(4E), SPD-5(4A) alone can also build a small amount of PCM that is weak but is prematurely dissembled by pulling forces. Furthermore, this observation supports the idea that PCM scaffold assembly and strength are interrelated. We suspect that the underlying molecular basis for overall PCM weakness is different for the 4E and 4A mutants (see Discussion). We conclude that dysregulated phosphorylation of SPD-5 leads to force-dependent and –independent defects in PCM assembly.

### 2.6 Proper SPD-5 phosphorylation is important for the fidelity of chromosome segregation

We hypothesized that PCM integrity defects caused by dysregulated phosphorylation of SPD-5 could interfere with centrosome function. To test this, we imaged chromosome segregation from metaphase until telophase in one-cell embryos solely expressing *gfp::spd-5* transgenes. Chromosome segregation was normal in all *spd-5(WT)* embryos. On the contrary, chromosome segregation was defective in 100% of *spd-5(4A)* and 14% of *spd-5(4E)* embryos (Figure 6A,B,D). Defects included failure to position mitotic chromosomes at the metaphase plate, lagging chromosomes during anaphase, and appearance of extranuclear DNA in telophase (Figure 6B; Figure S4A,B). 34% of *spd-5(4E)* embryos did not hatch (Figure S4C), and previous studies demonstrated that *spd-5(4A)* embryos are largely inviable (Woodruff et al., 2015; Wueseke et al., 2016). Thus, either hypo- and hyper-phosphorylation of SPD-5 interferes with chromosome segregation and embryonic development.

**Figure 6:**
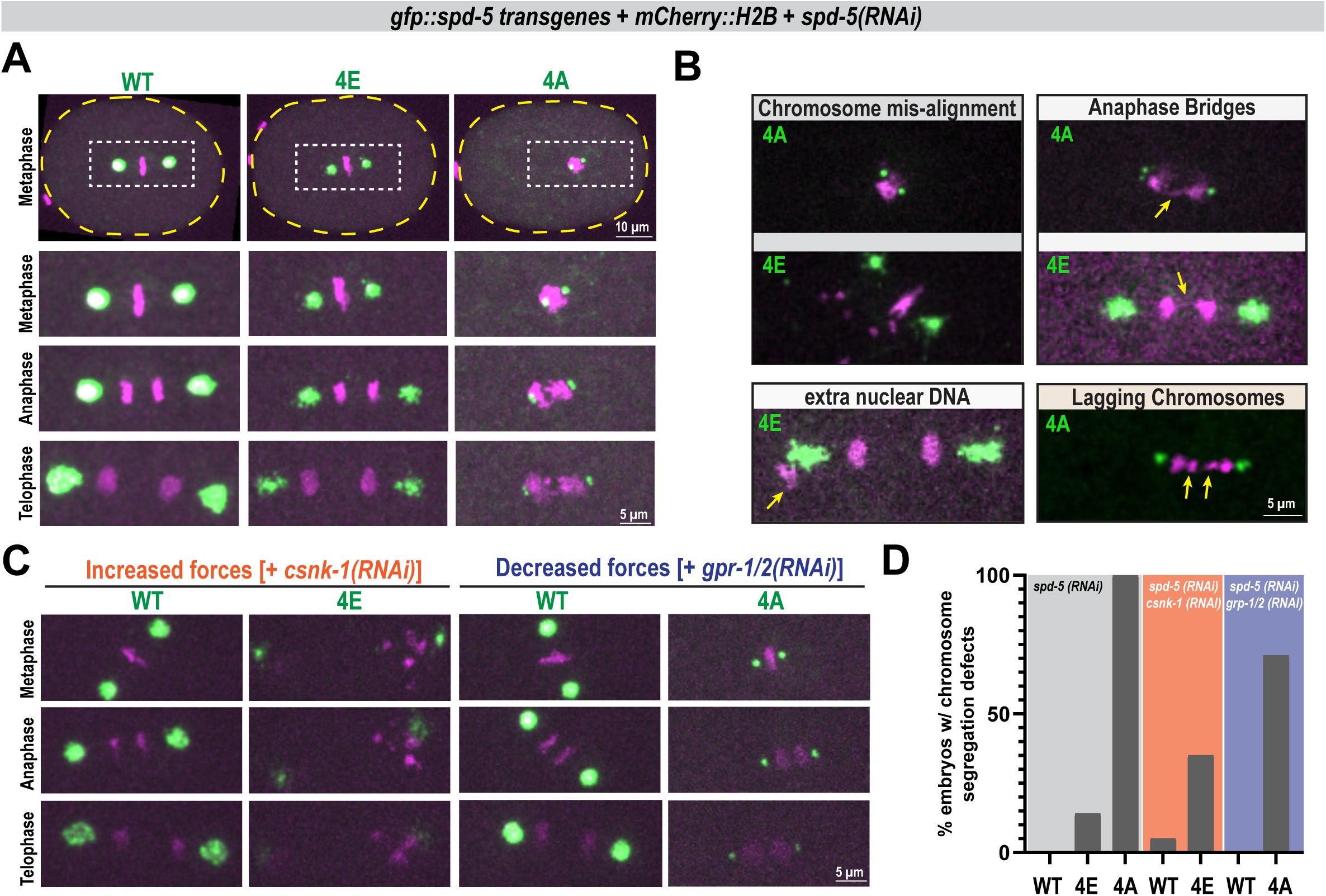
Proper SPD-5 phosphorylation is important for the fidelity of chromosome segregation. a. Representative images of chromosome segregation in one-cell embryos expressing *gfp::spd-5 transgenes* treated with *spd-5(RNAi)* to deplete endogenous SPD-5, imaged from metaphase until telophase. DNA is visualized with H2B::mCherry. b. Examples of chromosome segregation defects observed in (A). c. Chromosome segregation in transgenic *spd-5(RNAi)* embryos combined with *csnk-1(RNAi)* to increase microtubule pulling forces (Left) or *gpr-1/2(RNAi)* to reduce microtubule pulling forces (Right). d. Quantification of chromosome segregation defects in (A) and (C) reported as Percentage of embryos with chromosome segregation defects in (A) and (C). (gray panel: n= 29,29,23 (WT,4E,4A) embryos across 4 biological replicates; orange panel: n=20,17 (WT,4E) across 3 biological replicates; blue panel: (WT n=15 (WT,4A) across 3 biological replicates).

Our extrusion data indicated that PCM built solely from SPD-5(4E) is mechanically compromised, which could contribute to the mild chromosome segregation defect in these embryos. If true, sensitizing the system by increasing mechanical load should exacerbate PCM integrity and function defects in *spd-5(4E)* embryos. To test this, we increased cortically directed microtubule pulling forces ∼1.5-fold by depleting CSNK-1 (Panbianco et al., 2008). In *spd-5(WT)* + *csnk-1(RNAi)* embryos, spindle rocking was exaggerated, but PCM remained intact, as expected (Mittasch et al., 2020). In contrast, in *spd-5(4E)* + *csnk-1(RNAi)* embryos, PCM was violently ripped apart and caused an increase in chromosome segregation defects (Figure 6C,D; quantification of all defects is shown in Figure S4A,B). Since CSNK-1 depletion is known to impair polar body extrusion (Flynn and McNally, 2017), we excluded defects resulting from incorporation of the polar bodies into aberrant additional nuclei. These data further demonstrate that PCM built from SPD-5(4E) is mechanically unsound and cannot function in cases of increased load, unlike wild-type PCM.

We next tested if chromosome segregation defects seen in *spd-5(4A)* embryos were due to disrupted PCM mechanical integrity. Since chromosome segregation defects are already severe in *spd-5(4A)* embryos, it is not productive to further sensitize the system with *csnk-1(RNAi)*. Instead, we tested if decreasing microtubule pulling forces, via *gpr-1/2(RNAi)*(Colombo et al., 2003) could rescue chromosome segregation. We saw a 30% reduction in segregation errors for *spd-5(4A)* embryos with reduced forces, whereas chromosome segregation in wild-type embryos was not affected (Figure 6C,D). PCM size was still reduced in *spd-5(4A)* embryos and spindle centration and rocking were defective, indicating that the double RNAi knockdown was effective. We conclude that PCM built with SPD-5(4A) is too weak to resist physiological microtubule pulling forces, which ultimately affects the ability of PCM to assemble and function. Overall, our results demonstrate that the mechanical integrity of the PCM scaffold is important for its function in faithfully segregating mitotic chromosomes.

## 3.0 Discussion

The protein-rich PCM scaffold resists microtubule pulling forces needed to position the mitotic spindle and segregate mitotic chromosomes. Here, we investigated how the dynamics and material properties of the PCM scaffold influence its function in *C. elegans*. Our results support a model whereby PLK-1 phosphorylation of SPD-5 reduces its dynamics, which increases the viscoelasticity of PCM. Thus, PLK-1 phosphorylation promotes the initial assembly of the SPD-5 scaffold and its emergent material properties. Properly tuned viscoelasticity is required for the PCM to resist pulling forces and segregate chromosomes during mitosis.

Viscoelastic materials exhibit a hybrid response to stress: an elastic spring-like response that stores energy and a viscous response that dissipates energy over time. Our results indicate that the PCM scaffold is viscoelastic, a property that is increased by PLK-1 phosphorylation. Specifically, PLK-1 phosphorylation of the main scaffold protein SPD-5 decreased its dynamics within PCM condensates and concomitantly increased the viscous and elastic moduli of PCM condensates in vitro (Figure 1, 2). We saw similar results in vivo, where SPD-5 dynamics and PCM material properties changed in relation to phosphorylation status. Because both PLK-1 and its opposing phosphatase PP2A coexist within the PCM, SPD-5 likely exists in both phosphorylated and dephosphorylated states (Enos et al., 2018; Magescas et al., 2019). We propose that this balance tunes the viscoelastic properties of the PCM scaffold to optimize function (Figure 7). Under-phosphorylated SPD-5 is dynamic and can rearrange and depart, thus contributing to the overall viscous, liquid-like behavior. On the other hand, phosphorylated SPD-5 is less dynamic and cannot rearrange, thus contributing to the overall elastic, solid-like character of PCM. This viscoelastic character allows PCM to assemble properly and resist microtubule pulling forces during spindle assembly. Consistent with our model, dysregulation of the phosphorylation state of SPD-5 is pathological, leading to premature PCM disassembly, defective chromosome segregation, and embryonic lethality. PCM built with solely phospho-null SPD-5 (SPD-5(4A)) assembles a thin layer of dynamic PCM that is unable to resist microtubule pulling forces needed for full-scale assembly and proper chromosome segregation. On the other hand, PCM built from solely phospho-mimetic SPD-5 (SPD-5(4E)), exhibits a similar phenotype but is driven by a different mechanism. This PCM is less dynamic, exhibiting multiple hyperstable foci, which cannot cohere into a uniform PCM body and thus are broken apart by microtubule pulling forces. Our results suggest that SPD-5 dynamics reflect the functionally relevant viscoelastic properties of the PCM scaffold in embryos. Our *in vitro* data revealed that PLK-1 activity decreased SPD-5 dynamics and concomitantly increased the viscoelasticity of PCM condensates. *In vivo,* FRAP of PCM in metaphase-arrested cells showed SPD-5 turnover in matured PCM over a biologically relevant time scale (140 s). Thus, SPD-5 is dynamic, albeit at a low amount, consistent with molecular rearrangement expected for a viscoelastic material. Our group previously demonstrated that PCM can resist deformation induced by thermoviscous pumping at metaphase but not during anaphase or when PLK-1 is inhibited (Mittasch et al., 2020). These material states align with changes in SPD-5 dynamics we observed in the present study: SPD-5 became more dynamic in anaphase (Figure 4C-E) or when underphosphorylated (e.g., 4A mutant) (Figure 3C-D, 4F). Taken together these data support that SPD-5 dynamics are responsible for the viscoelastic properties of PCM in embryos.

**Figure 7:**
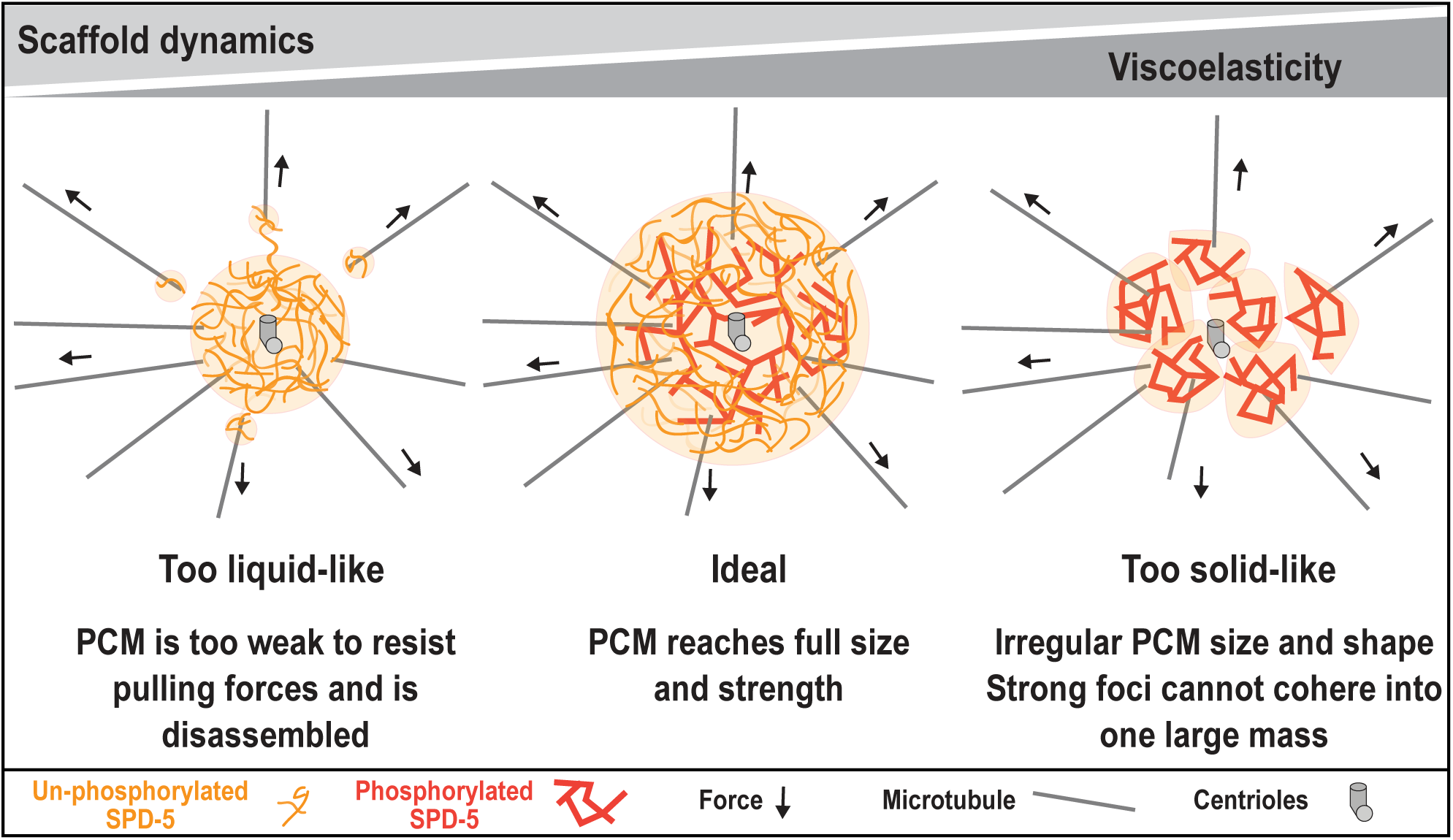
Phosphorylation tunes the functionally important viscoelastic properties of the PCM. Diagram of PCM architecture in a *C. elegans* embryo. Properly tuned PCM dynamics and viscoelasticity ensures PCM reaches full size and strength needed for function. Left, under-phosphorylated SPD-5 (orange) builds PCM that is too dynamic and weak, leading to a liquid-like PCM that gets quickly pulled apart by microtubule pulling forces. Right, over-phosphorylated SPD-5 (red) builds PCM that lacks dynamic character leading to solid-like PCM that is irregularly shaped and lacks mechanical integrity.

How important are these viscoelastic properties for function? PCM built from either SPD-5(4A) or SPD-5(4E) mutants disrupted PCM dynamics and assembly and caused chromosome segregation defects. A secondary question thus arises: is PCM scaffold size or dynamics most critical for function? A previous study revealed that embryos expressing a simpler SPD-5 phospho-mutant (S653A, S658A; called “2A”), failed to build full-sized PCM like SPD-5(4A)(Ohta et al., 2021). Partial photobleaching revealed that GFP::SPD-5(2A) and GFP::SPD-5(WT) both have similarly low dynamics, while GFP::SPD-5(4A) has high dynamics (Figure S5A,B), as shown in other experiments. However, PCM built from solely SPD-5(2A) properly segregates mitotic chromosomes (Ohta et al., 2021) while PCM built from solely SPD-5(4A) cannot. This suggests that achieving correct scaffold dynamics—and, by association, material properties—is more important than achieving full size for PCM function.

Why would PCM built from solely phospho-mimetic SPD-5, which we hypothesize is more viscoelastic than PCM built from phospho-null SPD-5, exhibit assembly defects? Since PLK-1 phosphorylation is essential for mitotic PCM assembly (Conduit et al., 2014; Decker et al., 2011; Haren et al., 2009; Lee and Rhee, 2011; Woodruff et al., 2015), it was surprising that hyper-phosphorylation of SPD-5 led to PCM that was smaller and irregularly shaped (Figures 3-6). Furthermore, this PCM existed as multiple foci unable to cohere into a uniform body, that then fragmented into many stable packets upon extrusion (Figure 4G) or under increased pulling forces (Figure 6). One hypothesis is that excessive phosphorylation increases the affinity of intermolecular SPD-5 interactions such that they become too strong and are unable to release and bind new molecules. Eventually, all possible interaction motifs per SPD-5 molecule would be occupied, thus preventing the addition of new molecules and capping growth. In theory, this would lead to high affinity clusters of SPD-5 that are weakly connected to each other. Simulations of ideal multivalent polymers demonstrated that this mechanism, termed “valency exhaustion”, is sufficient to cause kinetic arrest and stop condensate growth (Ranganathan and Shakhnovich, 2020). Consistent with this idea, we saw that PLK-1 phosphorylation initially promoted SPD-5 assembly but then inhibited growth and prevented PCM condensate fusion *in vitro* (Figure 1B,C). Alternatively, SPD-5 phosphorylation could create a new type of interaction between SPD-5 molecules, which then leads to new material properties. This is the case for amyloid formation in FUS condensates (Ranganathan and Shakhnovich, 2022). Our microrheology showed that unphosphorylated PCM condensates have one crossover between G’ and G" at low frequency. In phosphorylated PCM condensates, G’ and G" cross over at even lower frequency, indicating an increase in elastic character, but they also approach toward a second crossover point at higher frequency. A second crossover would indicate that the condensate has acquired more complex material properties, but what this means at the molecular level is still unclear (Liu et al., 2006). Regardless of the finer mechanism, our results argue that hyperphosphorylation increases intermolecular interactions between neighboring molecules at the expense of less connectivity at the mesoscale, thus leading to defects in overall PCM assembly and mechanical integrity.

The tunable viscoelastic nature of the PCM scaffold could help make sense of previous conflicting results concerning PCM dynamics and architecture. For example, *C. elegans* PCM is spherical and grows isotopically (Laos et al., 2015), which is expected for a viscous liquid. On the other hand, in situ cryo-electron tomography showed that PCM contains an underlying fibrous meshwork (Tollervey et al., 2025), which would be expected for an elastic solid. One mystery arising from this work was that the pore size of the meshwork was ∼6-8 nm, which, if static, would be too small to accommodate large PCM proteins and complexes (e.g., TPXL-1, ZYG-9, γ-tubulin ring complex) (Tollervey et al., 2025). A viscoelastic PCM could achieve these complex features. Viscous behavior over longer time scales would allow for rearrangements in the scaffold. This would allow pore dilation needed accommodate larger proteins and movement of material to achieve isotropic growth. Elastic behavior over longer time scales would maintain the overall structure of the scaffold and allow it to resist microtubule pulling forces.

Recently, it was shown that PCM in *D. melanogaster* embryos is a composite of a stable scaffold of Cnn proteins co-existing with a dynamic phase of TACC protein (Wong et al., 2025). The Cnn scaffold seems to be important for mechanical strength (Lucas and Raff, 2007), while the TACC scaffold is required to recruit specific clients (Wong et al., 2025). This combination of static and dynamic scaffolds suggests that fly PCM is also viscoelastic. Future work should use rheology to test if PCM from flies and other species is similarly viscoelastic and regulated by phosphorylation. We speculate that tunable viscoelasticity is a universal design principle for PCM across eukaryotes.

## ACKNOWLEDGEMENTS

We thank Jessica Feldman for advice on performing embryo laser extrusion, Jordan Raff for advice on *in vivo* experiments, and Anthony Hyman for antibodies.

## DATA AVAILABILITY

Further requests and information for resources and reagents should be directed to and will be fulfilled by the Lead Contact, Jeffrey Woodruff.

## AUTHOR CONTRIBUTIONS

M. Amato performed all experiments and analyzed data. J.H. Hwang performed the microrheology experiments. M.U. Rios developed and optimized the embryo extrusion assay and performed and analyzed extrusion experiments. N.E. Familiari made baculoviruses and expressed proteins. J.B. Woodruff made transgenic *C. elegans* strains. M. Amato and J.B. Woodruff wrote the manuscript. M.K. Rosen and J.B. Woodruff supervised the project.

## MATERIALS AND METHODS

### Experimental model and subject details

For the expression of recombinant proteins (listed in Table S1), we used SF9-ESF *S. frugiperda* insect cells grown at 27**°**C in ESF 921 Insect Cell Culture Medium (Expression Systems) supplemented with Fetal Bovine Serum (2% final concentration). *C. elegans* worm strains were grown on nematode growth media (NGM) plates at 23**°**C, following standard protocols (www.wormbook.org). Worm strains used in this study are listed in Table S2.

### Generation of transgenic *C. elegans*

*C. elegans* worm strains used in this study were created with MosSCI (Frokjaer-Jensen et al., 2008) and are based on constructs made previously (Woodruff et al., 2015). GFP::SPD-5(4E) mutant was created by replacing a section of the wild-type sequence with a mutated gene block. All plasmids were purified using a NucleoBond Xtra Midi Prep Kit (Macherey Nagel), combined with co-injection plasmids, and injected into strain EG6699 (tt5605, Chr II). After one week, worms were heat-shocked for 3 hr at 35℃ to kill worms maintaining extrachromosomal arrays. Moving worms without fluorescent co-injection markers were selected as candidates. Sequencing was used to confirm transgene integration.

### Protein purification

All expression plasmids are listed in Table S1. SPD-5, TPXL-1 and PLK-1 constructs were expressed and purified as previously described (Woodruff et al., 2017; Woodruff and Hyman, 2015; Woodruff et al., 2015). Baculoviruses were generated using the FlexiBAC system (Lemaitre et al., 2019) in SF9 cells. Protein was harvested 72 hr after infection during the P3 production phase. Cells were collected, washed, and resuspended in harvest buffer (25 mM HEPES, pH 7.4, 150 mM NaCl). All subsequent steps were performed at 4°C. Cell pellets were resuspended in buffer A (25 mM HEPES, pH 7.4, 30 mM imidazole, 500 mM KCl, 0.5 mM DTT, 1% glycerol, 0.1% CHAPS) + protease inhibitors and then lysed using a dounce homogenizer. Proteins were bound to Ni-NTA (Qiagen), washed with 10 column volumes of buffer A, and eluted with 250 mM imidazole. For SPD-5 proteins, Ni-NTA eluate was then bound to MBP-Trap beads (Chromotek) and washed with 5 column volumes of buffer A and eluted with by adding PreScission protease, incubating overnight and then passing over Ni-NTA to remove the Precission protease. For TPXL-1 proteins, Ni-NTA eluate was equilibrated with binding buffer (25 mM HEPES, pH 7.4, 150 mM KCl) and then bound to SP Sepharose (GE Healthcare catalog No: 10261262) and washed with 5 column volumes of binding buffer. TPXL-1 was eluted using buffer A. For PLK-1 proteins, Ni-NTA eluate was concentrated with a 30 K MWCO Amicon concentrator (Millipore), filtered, and then passed over a Superdex 75 increase size exclusion column (Cytiva). Purified proteins were then concentrated using 30 K MWCO Amicon concentrators (Millipore). All proteins were aliquoted in PCR tubes, flash-frozen in liquid nitrogen, and stored at −80°C. Protein concentration was determined by measuring absorbance at 280 nm using a NanoDrop ND-1000 spectrophotometer (Thermo Fisher Scientific).

### RNAi treatment

RNAi was done by feeding. RNAi feeding using *gpr-1/2*, *csnk-1*, feeding clones from the Ahringer and Vidal collections (Source BioScience;(Rual et al., 2004)) The *spd-5* feeding clone targets a region that is reencoded in our MosSCI transgenes (Woodruff et al., 2015). Bacteria were seeded onto nematode growth media (NGM) supplemented with 1 mM isopropyl β-D-1-thiogalactopyranoside (IPTG) and 100 µg/mL ampicillin. For chromosome segregation experiments, *spd-5* feeding clone was combined with either *csnk-1* or *gpr-1/2* feeding clone and plated as before. L4 hermaphrodites were grown on feeding plates at 23°C for 24-34 hr or at 16°C for 24-34 hr in worms expressing *emb-30(ts)*.

Target sequence for *spd-5(RNAi)*

5′-TGG AAT TGT CCG CTA CTG ATG CAA ACA ACA CAA CTG TCG GAT CTT TTC GTG GAA CTC TTG ATG ACA TTC TGA AGA AAA ACG ATC CAG ATT TCA CAT TAA CCT CTG GTT ATG AAG AAA GAA AGA TCA ACG ACC TGG AGG CAA AGC TCC TCT CTG AGA TCG ACA AGG TAG CTG AGC TGG AAG ATC ACA TTC AGC AGC TCC GTC AAG AAC TTG ACG ACC AAT CTG CAA GGC TTG CCG ATT CAG AAA ATG TTC GCG CTC AGC TTG AAG CGG CCA CTG GAC AAG GAA TCC TCG GAG CTG CTG GAA ACG CTA TGG TTC CAA ATT CAA CGT TCA TGA TCG GGA ACG GTC GTG AAT CAC AGA CGC GAG ACC AGC TCA ATT ACA TTG ATG ATC TTG AAA CGA AGT TAG CTG ATG CGA AGA AGG AAA ATG ATA AGG CTC GTC AGG CAC TCG TTG AAT ACA-3′.

### Western blotting

45 adult worms were picked and transferred to blank plates for 20 min to remove bacteria from their bodies. Worms were then moved to PCR tubes containing 10 µl of milli-Q water to which 10 µl of SDS loading buffer was added. Samples were separated by SDS-PAGE. Protein from each gel was transferred to a nitrocellulose membrane using a Trans-blot turbo transfer for high molecular weight proteins (10 min). Membranes were incubated in a blocking buffer consisting of TBS-T + 3% Blotting-Grade Blocker (BioRad) shaking at room temperature for 1 hr. Membranes were then washed three times with TBS-T and incubated shaking with primary antibodies overnight at 4°C. The primary antibody was washed off three times with TBS-T and incubated with secondary antibodies at room temperature for 1 hr. Each membrane was then incubated in ECL reagent (Thermo Fisher Scientific SuperSignal West Femto) for 5 min and imaged with a ChemiDoc Touch Imaging System. Primary antibodies: Mouse anti-alpha tubulin (Product No: 3873S; 1:1,000; Cell Signaling Technologies); Rabbit anti-SPD-5 C-terminus (1:1,000, clone 785, Dresden Antibody Facility; Pelletier et al., 2004). Secondary Antibodies (1:50,000 for all): HRP conjugated Goat anti-Rabbit IgG (1 mg/ml) (catalog No: 65-6120, catalog No: J276300; Invitrogen); HRP conjugated Goat anti-Mouse (1.5 mg/ml) (Catalog No: 62-6520, Catalog No: WA312227; Invitrogen); and HRP conjugated Donkey anti-Goat (1 mg/ml) (Catalog No: A15999, Catalog No: 58-155-072318; Invitrogen).

### Microscopy of *C. elegans* embryos

Embryos from adult worms were dissected on a 22 × 22 mm coverslip (Catalog No: 2975-225; Coring) using a 22-gauge needles in a 7 µl solution of egg salts buffer (ESB) (118mM NaCl, 48mM KCl, 2mM CaCl_2_, 2mM MgCl_2_, 25mM Hepes) or M9 buffer. Samples were then mounted onto plain 25 × 75 × 1 mm microscope slides (Catalog No: 12-544-4; Fisher Scientific). Time-lapse images were acquired with an inverted Nikon Eclipse Ti2-E microscope with a Yokogawa confocal scanner unit (CSU-W1), piezo Z stage, and an iXon Ultra 888 EMCCD camera (Andor), controlled by Nikon Elements software. We used a 100X Plan Apo silicon immersion objective (NA 1.35). To quantify GFP::SPD-5 transgene intensity we imaged at 15-18 × 1-µm Z-stacks using 488-nm at 2 x 2 binning, 15% laser power, 100 ms exposures every 20 seconds until centrosome disassembly except where specified.

For microtubule depolymerization assays, one-cell embryos from adult worms were treated with ESB mixed with nocodazole to make a 40 µM nocodazole solution or 2% DMSO solution and imaged prior to PNM. For SPD-5 growth curves one-cell embryos from adult worms were dissected in M9 and imaged prior to pronuclear meeting (PNM).

### *In vivo* FRAP of metaphase arrested embryos

Metaphase arrest was achieved by temperature inactivation of *emb-30(ts)* by incubating embryos from 16℃ to 25℃ for 15 minutes. One-cell embryos from adult worms were dissected in M9 buffer and identified prior to metaphase. Metaphase arrest was determined by chromosomes positioned at the metaphase plate. After 200 s post metaphase arrest, stable PCM was imaged before photobleaching and every 20 s after photobleaching for 140 s. PCM was photobleached with a 405-nm laser using a roi the area of the centrosome at 20% intensity, 250 µs dwell time. Mean GFP::SPD-5 transgene intensity was measured and normalized to the mean intensity of the centrosome prior to photobleaching as the highest value and cytoplasmic region as the lowest value. To validate PCM mass was stable during arrest, the mean GFP::SPD-5 transgene intensity of the unbleached centrosome was recorded.

### *In vivo* partial FRAP of centrosomes

One-cell embryos from adult worms were dissected in M9 buffer and photobleached with a 405-nm laser at NEBD using a stim point positioned in the center of centrosomes at 1% intensity, 50 ms dwell time. To quantify the intensity of GFP::SPD-5 transgene after photobleaching a 5 µm line scan was positioned at centrosomes such that the maximum and minimum intensity occurred at 1.82 µm and 2.6 µm respectively. GFP::SPD-5 transgene intensity was recorded immediately after photobleaching and 100 s thereafter. Intensities are averaged over replicates and normalized to the maximum intensity immediately after photobleaching.

### Centrosome extrusion assay

Dissected one-cell embryos from adult worms were coated in a solution of 10 µm Calcofluor-white (Biotium catalog No: 29067) in high salt buffer (150mM KCl, 25mM Hepes pH 7.4). or MQ H20 with 25mM Hepes pH 7.4. In PP2A inhibition experiments, centrosomes were extruded into high salt buffer containing 10 µM LB-100. To extrude centrosomes, a 405 nm laser was targeted to the periphery of the eggshell using (100% laser intensity, 500 ms dwell time). Cell cycle stage was determined by chromosome organization via mCherry::H2B signal. Imaging was performed as above with a 60 × 1.2 NA Plan Apochromat water-immersion objective, GFP/mCherry splitter, 2 x 2 binning, 488 nm laser (15% intensity), 561 nm laser (10% intensity), 51 x 0.5 µm Z-steps, every 2 min for 60 min or until centrosomes completely dissolved. The integrated density of GFP::SPD-5 was normalized to centrosome intensity prior to extrusion in the normalized plot.

High resolution imaging of GFP-SPD-5(4E) was performed using 100X Plan Apo silicon immersion objective (NA 1.35) imaged at 15-18 × 1-µm Z-stacks using 488-nm at 2 x 2 binning, 15% laser power, 100 ms exposures every 5 s until centrosomes left the field of view.

### Chromosome segregation defects

To quantify chromosome segregation defects in embryos expressing *gfp::spd-5 transgenes* treated with various RNAi one-cell embryos were dissected into M9 and imaged prior to metaphase until telophase. Chromosome segregation defects were determined by organization of DNA visualized by mCherry::H2B signal. Embryos were imaged at 15-18 × 1-µm Z-stacks using 488-nm at 2 x 2 binning, 15% laser power, 561-nm at 2 x 2 binning, 20% laser power, 100 ms exposures every 20 seconds.

### 96-well glass bottom passivation

To clean a 96-well glass bottom imaging plate, wells were then treated with a solution of 5% v/v Hellmanex (Sigma-Aldrich catalog No: Z805939) for 1hr and washed 10X in H2O and dried. A solution of 5M NaOH is added to wells for 1hr and washed 5X with H2O. To passivate the surface of the wells, the 96-well glass-bottom imaging plate was treated with a solution of 0.5% w/v of mPEG-Silane, MW 5,000 (Creative PEGWorks catalog No: PLS-2011) in 95% EtOH for 1hr and then washed 10X with H2O and dried. On the day of the experiment, a solution of 5% w/v of Bovine Serum Albumin (BSA) (Sigma-Aldrich) was incubated in the wells for 30 min, washed 5X with H2O and then dried.

### *In vitro* PCM scaffold reconstitution and imaging

To reconstitute PCM scaffold assembly from purified components we incubated 1µM SPD-5::RFP, 1µM TPXL-1::GFP, 75mM KCl, 1mM DTT, 0.4mM ATP, 1mM MgCl_2_, 4mM Hepes pH 7.4, except where indicated, in a passivated 96-well glass-bottom imaging plate for 15 min. To prevent evaporation wells are covered with parafilm (Bemis). To image reconstituted PCM we used at 100x silicon objective an inverted Nikon Eclipse Ti2-E microscope with a Yokogawa confocal scanner unit (CSU-W1), piezo Z stage, and an iXon Ultra 888 EMCCD camera (Andor), controlled by Nikon Elements software. Imaging was performed at 488-nm; 2 x 2 binning, 1% intensity, 100ms exposure, and 561-nm; 1x1 binning, 5% intensity, 100ms exposure at a single imaging plane at the surface of the glass except where specified. For condensate FRAP and fusion assays, we imaged at 0.5 x 10µm relative to the surface of the glass well.

### *In vitro* PCM scaffold FRAP assay

To measure the dynamics of reconstituted PCM scaffolds, we incubated samples (as above) with 0.5 µM PLK-1(CA) or 0.5 µM PLK-1(KD) for 30 minutes and then photobleached with a 405 nm laser using a rectangular ROI positioned within condensates at 20% intensity, 200 µs dwell time. We imaged condensates at 0.5 x 10 µm relative to the surface of the glass well. The mean intensity of SPD-5::RFP and TPXL-1::GFP was recorded prior to photobleaching and recorded every 15 seconds for 10 min post photobleaching. Mean intensities were normalized to mean intensity prior to photobleaching. 6 and 5 replicated were performed for PLK-1(KD) and PLK-1(CA) conditions respectively.

### *In vitro* PCM scaffold dilution assay

To determine the sensitivity to dilution of reconstituted PCM, we prepared reconstituted PCM scaffolds using 1µM SPD-5::RFP, 1µM TPXL-1::GFP, 50mM KCl, 1mM DTT, 0.4mM ATP, 1mM MgCl_2_, 4mM Hepes pH 7.4 and incubated with 0.5 µM PLK-1(CA) or 0.5 µM PLK-1(KD) in a cleaned non-passivated 96-well glass imaging plate. Samples were incubated for 1hr and Imaged. Samples were then diluted samples with 4.3x extrusion buffer (150mM KCl, 25mM Hepes pH 7.4) and incubated for an additional hour and imaged. We measured the total integrated density of SPD-5::RFP prior and after dilution in which 6 images at different locations at the surface of the glass well were gathered for each condition.

In the case of SPD-5(4E)::GFP, PCM condensates were prepared using 1µM SPD-5(4E)::GFP, 1µM TPXL-1, 50mM KCl, 1mM DTT, 4mM Hepes pH 7.4 n a cleaned non-passivated 96-well glass imaging plate. Samples were incubated and imaged as before but instead diluted 4.3x into (250mM KCl, 25mM Hepes pH 7.4).

### *In vitro* PCM scaffold assembly assay

To determine the effect of PLK-1 phosphorylation on the assembly of reconstituted PCM scaffolds, we incubated 1µM SPD-5::GFP, 1µM TPXL-1, 50mM KCl, 1mM DTT, 0.4mM ATP, 1mM MgCl_2_, 4mM Hepes pH 7.4 with 0.5µM PLK-1(CA) or 0.5µM PLK-1(KD) in a passivated 96-well imaging plate. We imaged using 488-nm; 2 x 2 binning, 1% intensity, 100ms exposure. 3 images at distinct locations at the surface of the glass well were recorded at every interval for intervals 5, 10, 15, 20, 30, 45, 60, 90 minutes from one experiment for PLK-1(CA) and PLK-1(KD). Condensate size was measured using a semi-automated threshold-based analysis. The watershed function in FIJI was used to segregated chains of condensates.

### Video particle tracking microrheology sample preparation

Samples were prepared in 384-well glass bottom microwell plates (Brooks Life Science Systems: MGB101-1-2-LK-L). Prior to use, the plates were cleaned with 5% Hellmanex III (Hëlma Analytics), etched with 1M KOH, and siliconized with Sigmacote (Sigma-Aldrich). On the day of the experiment, individual wells were blocked with 1% bovine serum albumin, then rinsed thoroughly with MilliQ-water. PCM samples were prepared with 1 μM SPD-5::GFP, 1 μM TPXL-1, 50 mM KCl, 0.4 mM ATP, 1 mM MgCl_2_, 1mM Hepes pH 7.4 pH), 150 beads/μL with 0.2 μM PLK-1(KD) or 0.2 μM PLK-1(CA) and 100 nm-diameter carboxylate-modified fluorescent beads (Invitrogen: F8801). The final concentration of beads was <150 beads/μL. Samples were incubated inside the temperature-controlled microscope chamber at 30°C for at least 1 hour before imaging.

### Video particle tracking microrheology microscopy

Images were captured on a Lecia DMI6000 B microscope base with a Yokogawa CSU-X1 spinning disk confocal scanner unit and a 405/488/561/647 nm Laser Quad Band Set filter cube (Chroma) with a plan apo 63 or 100 × 1.40 NA oil immersion objective. Images were acquired using a Hamamatsu ImagEMX2 EM-CCD camera at 15ms/frame using stream acquisition function on Metamorph (Biovision) software. At least 10000 frames (3-25 min) were acquired for each acquisition, up to 6 acquisitions were made per sample per session, and at least 3 sessions were carried out per sample for reproducibility.

### Video particle tracking microrheology data analysis

Particle tracking and calculation of mean squared displacement (MSD) was performed using MATLAB codes by Daniel Blair & Eric Dufresne (https://site.physics.georgetown.edu/matlab/code.html). Average MSD was calculated from >17000 individual trajectories and smoothed using a moving average with span <10% of total number of frames. Elastic (G’) and viscous (G”) moduli as a function of frequency (ω) were calculated from the averaged MSD from 0.015 to 100s using generalized Stokes-Einstein relation (GSER) as described by Mason TG (Rheologica Acta, 2000) using MATLAB codes by Andrew Sun (https://github.com/andrewx101/track_analysis/releases/tag/v2.05). Viscosity (η) is calculated (η = G”/ω) and plotted against frequency. From viscosity plot, the mean value of the plateau at the low frequency was used to estimate the zero-shear viscosity (η_0_).

### Image quantification and statistical analysis

Images were analyzed using a semiautomated, threshold-based particle analysis in FIJI (https://fiji.sc/). MATLAB (Mathworks) was used to analyze VPT experiments. All data are expressed as the mean ± 95% confidence intervals as stated in the figure legends and results. The value of n and what n represents (e.g., number of centrosomes, condensates or experimental replicates) is stated in figure legends and results. A Brown-Forsythe and Welch ANOVA statistical test were used for normally distributed data, and a Kruskal-Walis were used for non-normally distributed data. Significance was reported as; p value < 0.05 = *, p value < 0.005 = **, p value < 0.0005 = ***, p value of < 0.00005 = ****.

---

**TABLE S1.** Baculoviral expression constructs for protein expression.

| protein | sequence | plasmid name | parent plasmid (pOCC #) | N-term tag | C-term tag | species |
| --- | --- | --- | --- | --- | --- | --- |
| SPD-5 | Full-length | JWV1 | pOCC28 | MBP-PreScission | eGFP-PreScission-6xHis | *C.elegans* |
| SPD-5 | Full-length | JWV3 | pOCC25 | MBP-PreSci ssion | RFP-PreScis sion-6xHis | *C.elegans* |
| SPD-5 | Full-length | JWV10 | pOCC27 | MBP-PreSci ssion | eGFP-PreScis sion-6xHis | *C.elegans* |
| TPXL-1 | Full-length | JWV29 | pOCC27 | MBP-PreSci ssion | eGFP-PreSci ssi on-6xHis | *C.elegans* |
| TPXL-1 | Full-length | JWV30 | pOCC28 | MBP-PreSci ssion | PreScis sion-6xHis | *C.elegans* |
| PLK-1(CA) | Full-length T194D | JWV11 | pOCC7 | 6xHis-PreScis sion | - | *C.elegans* |
| PLK-1(KD) | Full-length K67M | JWV12 | pOCC7 | 6xHis-PreScis sion | - | *C.elegans* |

**TABLE S2.**
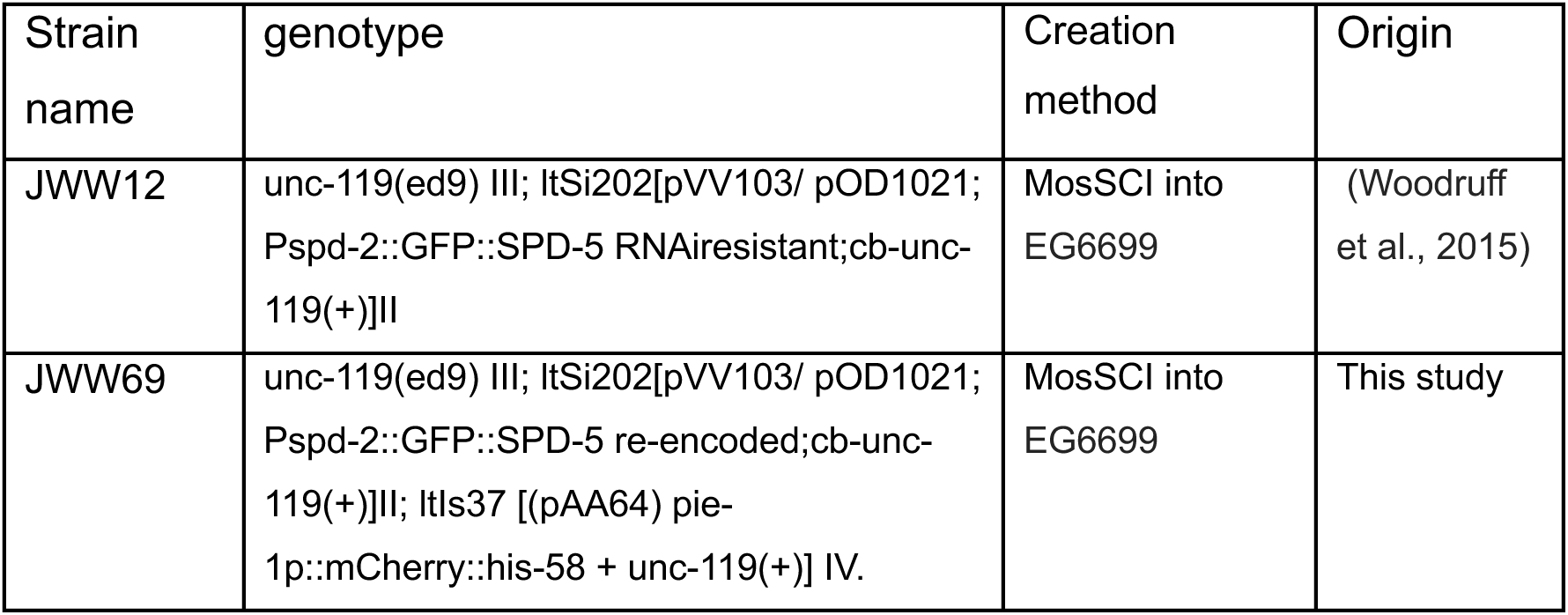
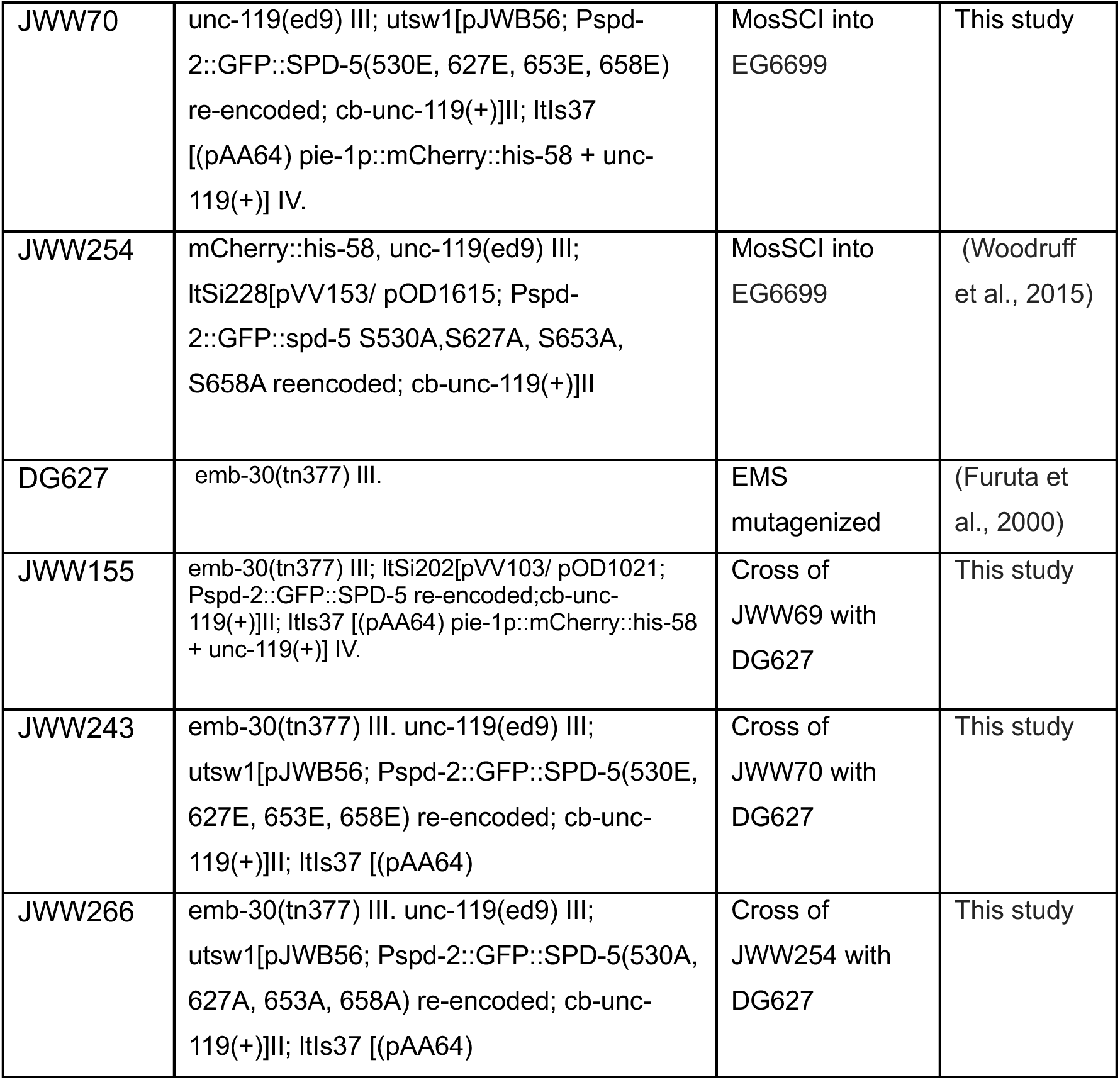
C. elegans strains.

## SUPPLEMENTAL FIGURE LEGENDS

**Figure S1.**
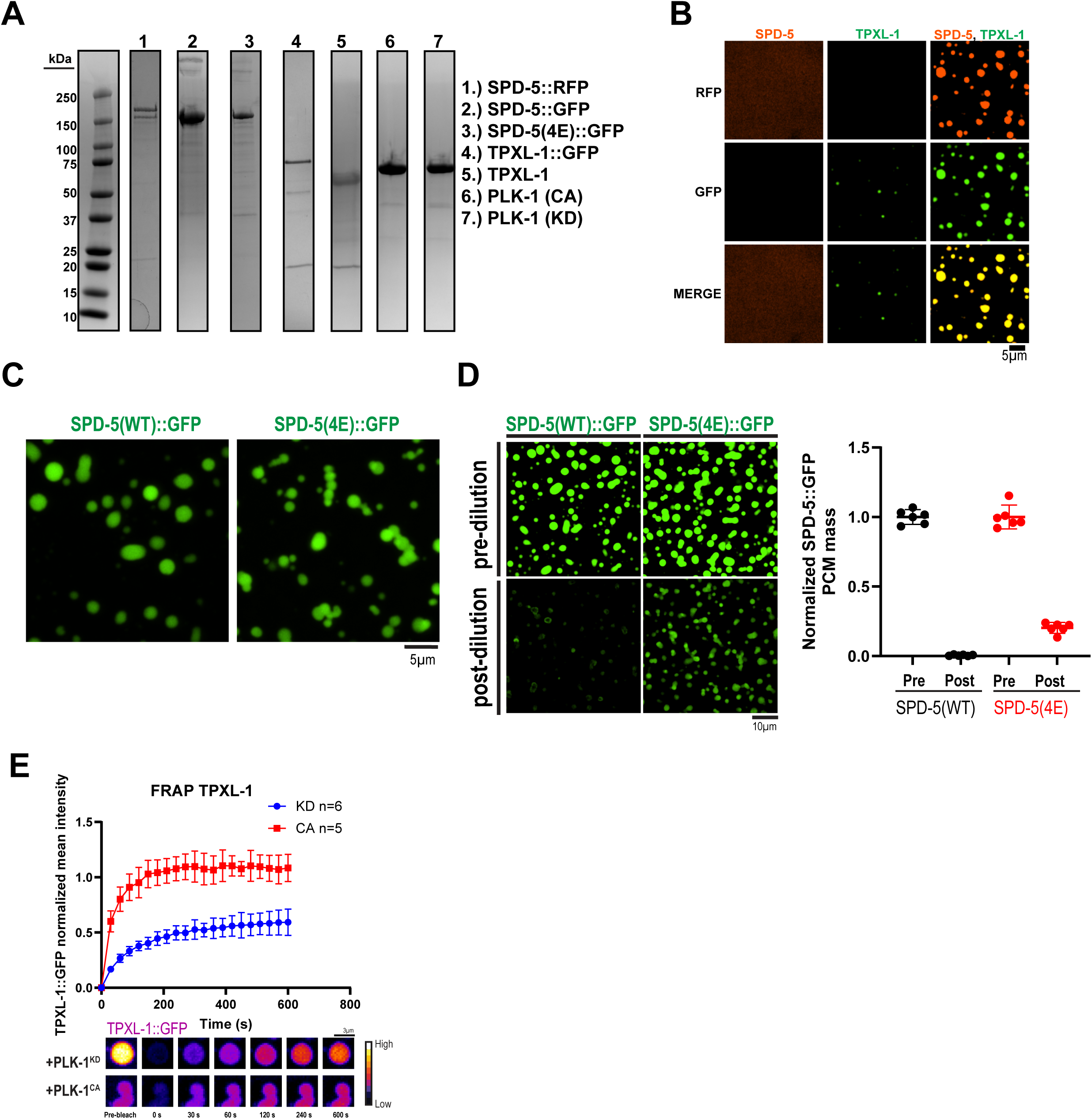
In vitro analysis of PCM. a. Coomassie gel of proteins used in this study b. Reconstituted PCM condensates. Representative images of reconstituted PCM condensates prepared with 1 µM SPD-5::RFP, 1 µM TPXL-1::GFP, 50 mM KCl, Hepes pH 7.4 and incubated for 15 min. c. Representative images of reconstituted PCM condensates prepared with either 1 µM SPD-5:GFP(WT) or 1 µM SPD-5(4E), 1 µM TPXL-1::GFP, 50 mM KCl, Hepes pH 7.4 and incubated for 30 min. d. Dilution assay of reconstituted PCM. 1 µM SPD-5::GFP(WT) or 1 µM SPD-5::GFP(4E), 1 µM TPXL-1::GFP was incubated in buffer (50 mM KCl, 4 mM Hepes pH 7.4, 10 mM DTT, 0.4 mM ATP, 1 mM MgCl_2_) with 0.5 µM PLK-1 (KD) or 0.5 µM PLK-1(CA) for 1 hr and imaged. Samples were then diluted 4.3X into buffer (200mM KCl, 25mM Hepes pH.7.4) and imaged after 1 hr (left panels). Right, quantification of SPD-5::RFP integrated density before and after dilution (mean +/− 95% C.I.; PLK-1(KD) n=6 images, PLK-1 (CA) n=6 images with >100 condensates). e. Fluorescence recovery after photobleaching (FRAP) of reconstituted PCM condensates incubated with 0.5 µM PLK-1(KD) or 0.5 µM PLK-1(CA) for 30 min. TPXL-1::GFP intensity was measured and normalized (mean +/− 95% C.I.; KD, n = 5, CA n = 6 condensates).

**Figure S2.**
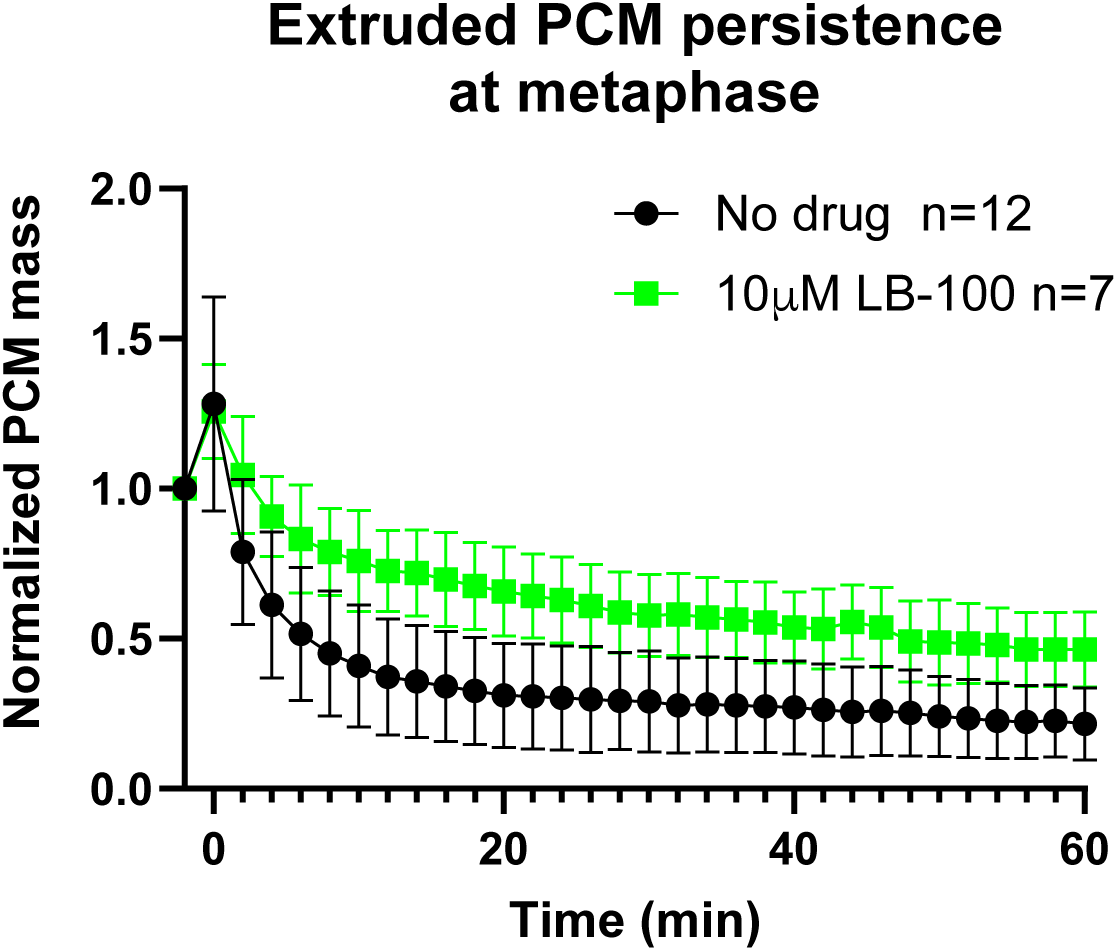
Effect of PP2A inhibition of PP2A on SPD-5 persistence in extruded PCM. Quantification of PCM from metaphase embryos expressing GFP::SPD-5(WT) + endogenous SPD-5 extruded into high salt buffer (150mM KCl, 25mM Hepes pH 7.4) (black curve), or high salt buffer with 10 µM LB-100 (green curve)(mean +/− 95% C.I.; no drug n= 12, 10 µM LB-100 n=7 centrosomes).

**Figure S3.**
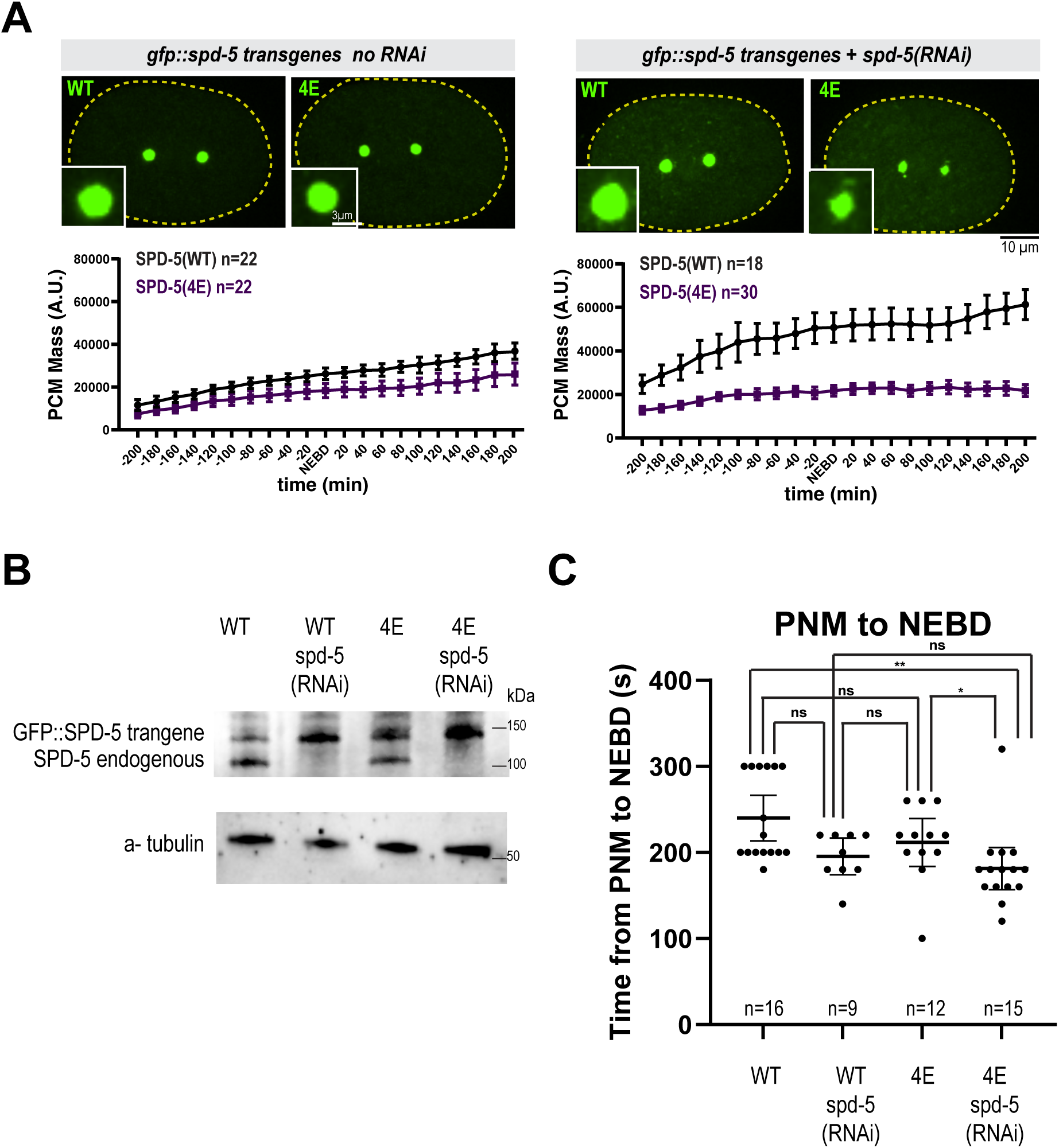
*In vivo* characterization of SPD-5(4E). a. PCM assembly in one-cell embryos expressing GFP::SPD-5 transgenes. Top left, representative images taken at nuclear envelope breakdown (NEBD). Bottom left, quantification of GFP::SPD-5 integrated density relative to NEBD. Top right, worms were treated with RNAi to deplete endogenous SPD-5 (mean +/− 95% C.I.; WT n=22, 4E n=22, WT + *spd-5(RNAi)* n=18, 4E + *spd-5(RNAi)* n=30). b. Western blot against SPD-5. Alpha tubulin was detected as a loading control. c. Time from pronuclear meeting (PNM) to nuclear envelope breakdown (NEBD) was measured in one-cell embryos expressing GFP::SPD-5 transgenes (mean +/− 95% C.I., p values from Kruskal-Wallis test followed by Dunn’s multiple comparisons test).

**Figure S4.**
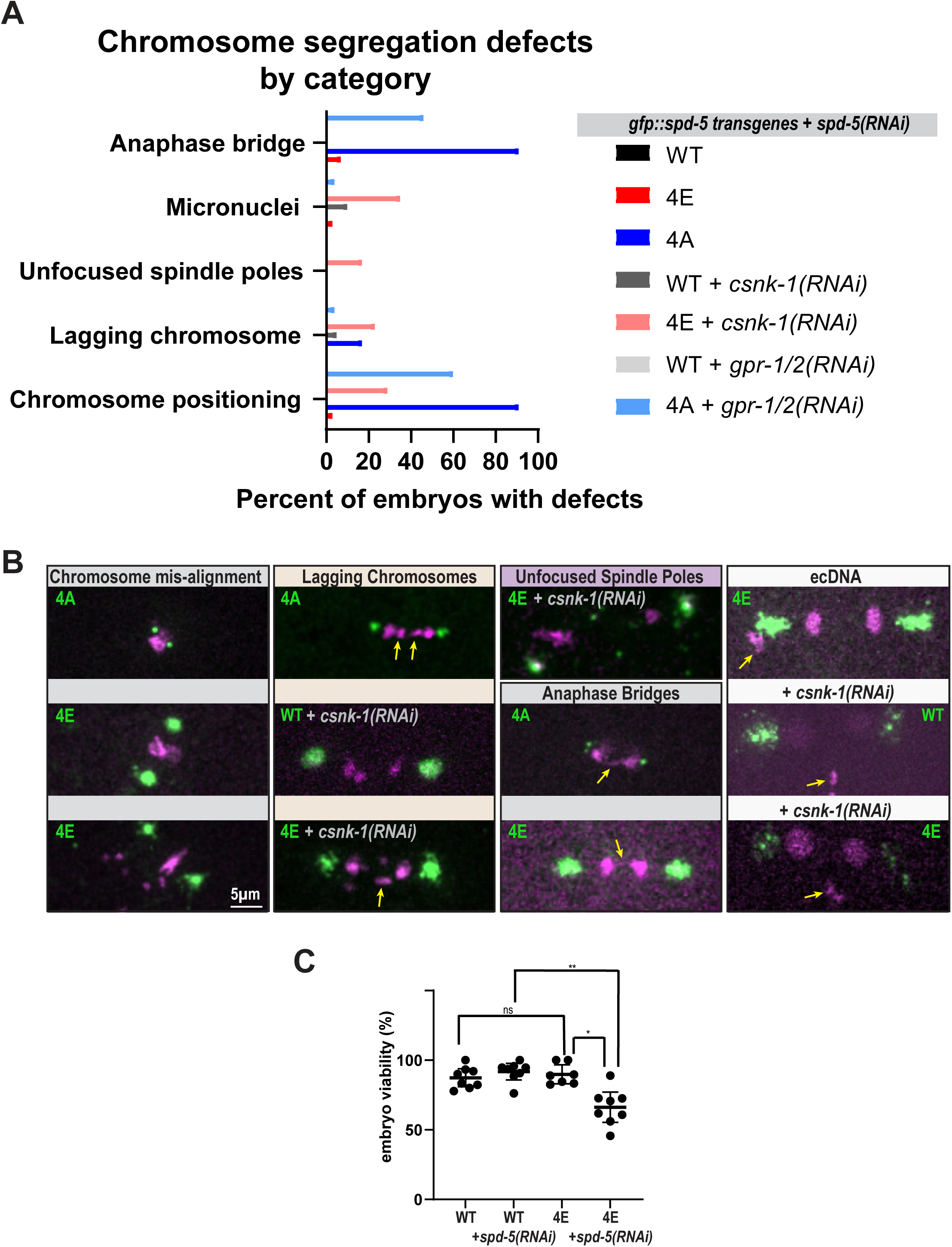
Characterization of chromosome segregation defects in embryos expressing *gfp::spd-5 transgenes*. a. Quantification of chromosome segregation defects by category in Figure 6A and C (see figure 6D for n values). b. Representative images of chromosome segregation defect categories. Unfocused spindle poles were observed in cases where centrosomes broke symmetry along the spindle axis. c. Embryo viability assay (mean +/− 95% C.I.; WT n=8, WT + *spd-5(RNAi)* n=8, 4E n=7, 4E + *spd-5(RNAi)* n=8 mothers, >20 embryos counted per mother; p values from a Kruskal-Wallis test followed by Dunn’s multiple comparisons test).

**Figure S5.**
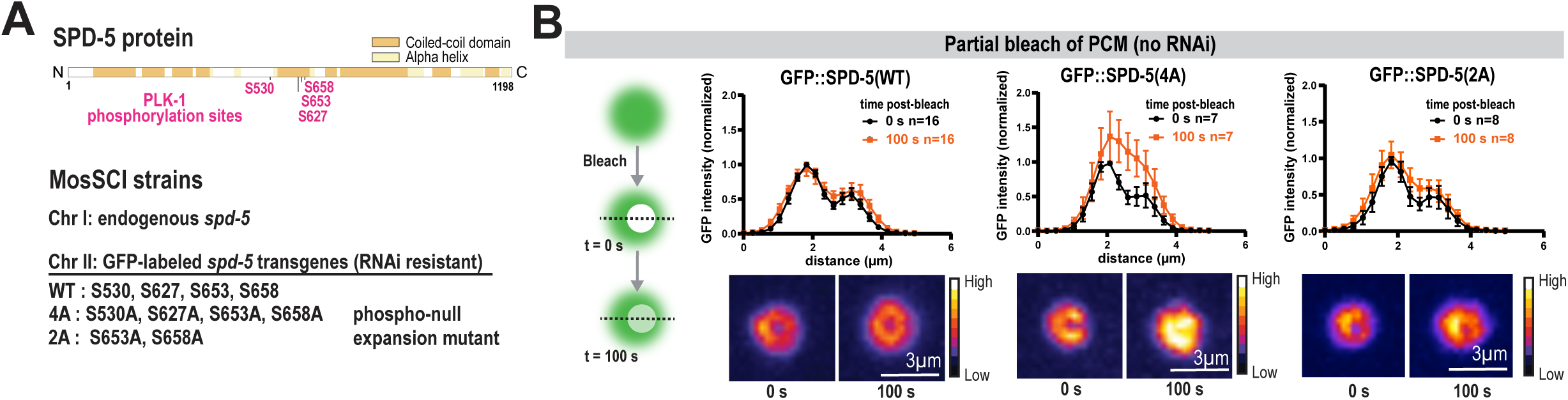
Partial FRAP of PCM in embryos expressing *gfp::spd-5* transgenes during mitosis. a. Top, diagram of key PLK-1 phosphorylation sites in SPD-5. Bottom, design of *gfp::spd-5* transgenes expressed at the Mos locus on chromosome II. GFP::SPD-5(2A) harbors two of the four S-to-A mutations present in GFP::SPD-5(4A). b. Partial fluorescence recovery after photobleaching (FRAP) of PCM at nuclear envelop breakdown in one-cell embryos (no RNAi). GFP intensity was measured with a 5 μm line scan 0 s and 100 s after photobleaching (mean +/− 95% C.I.; GFP::SPD-5(WT) n=16, GFP::SPD-5(4A) n=7, GFP::SPD-5(2A) n=8 centrosomes).

